# Modulation of Extracellular Matrix Composition and Chronic Inflammation by Pirfenidone Promotes Scar Reduction in Retinal Wound Repair

**DOI:** 10.1101/2023.11.22.538088

**Authors:** Laura Jahnke, Virginie Perrenoud, Souska Zandi, Yuebing Li, Federica Maria Conedera, Volker Enzmann

## Abstract

Wound repair in the retina is a complex mechanism and a deeper understanding of it is necessary for the development of effective treatments to slow down or even prevent degenerative processes leading to photoreceptor loss. In this study, we harnessed a laser-induced retinal degeneration model, enabling a profound molecular elucidation and a comprehensive, prolonged observation of the wound healing sequence in a murine laser-induced degeneration model until day 49 post laser. Our observations included the expression of specific extracellular matrix proteins and myofibroblast activity, along with an analysis of gene expression related to extracellular matrix and adhesion molecules through RNA measurements. Furthermore, the administration of pirfenidone, an anti-inflammatory and anti-fibrotic compound, was used to modulate the scar formation after laser treatment. Our data revealed upregulated collagen expression in late regenerative phases and sustained inflammation in the damaged tissue. Notably, treatment with pirfenidone was found to mitigate scar tissue formation, effectively downregulating collagen production and diminishing the presence of inflammatory markers. However, it did not lead to the regeneration of the photoreceptor layer.

## Introduction

Retinal fibrosis, which is the final phase of photoreceptor loss in retinal diseases such as in age-related macular degeneration (AMD), includes the formation of degenerated tissue in the eye as a result of inflammatory responses and wound healing. The wound healing process is typically segmented into four distinct phases: hemostasis, inflammation, proliferation, and maturation. Fibrosis is a pathological condition characterized by the excessive accumulation of fibrous connective tissue, primarily composed of collagen. This process is a response to chronic inflammation, injury, or other underlying disorders and can lead to impaired tissue function and structural alterations. Thereby, a fibrotic scar is formed by new extracellular matrix (ECM) hindering repair and resulting in vision loss [1,2]. In AMD, alteration of Bruch’s membrane and chronic inflammation may lead to photoreceptor loss [3]. Numerous publications have established a connection between increased cytokine expression and progression of AMD [4–7]. Furthermore, local inflammation resulting from the buildup of cellular debris has been found to trigger drusen formation in AMD patients across all stages [8]. IL-1β has been shown to cause neurodegenerative damage, particularly in tissue with long-term exposure [9,10]. Tumor necrosis factor–alpha (TNF-α), among others, has also been associated with the pathogenesis of AMD and increased expression of this factor was found in a laser-induced choroidal neovascularization (CNV) model in mice [11]. Re-inflammation during the wound healing process leads to an inhibition of regeneration as new healthy tissue is damaged and cytokines have a profound modulatory effect on the regenerative activity of endogenous stem cells and progenitor cells [12]. As a result of chronic inflammation, the axons of neurons in the affected area continue to lose their ability to interact with other surrounding neurons and thus lose their functionality to process visual information to the brain [13].

Therefore, we aim to inhibit fibrosis and inflammation during the proliferative stage of wound healing by studying the influence of the non-peptide synthetic molecule pirfenidone (PFD) on retinal degeneration after laser-induced damage in the outer nuclear layer (ONL). PFD is thereby blocking transforming growth factor-beta (TGF-β). TGF-β is a multifunctional factor in fibrotic remodeling, capable of triggering myofibroblast transition, activating Smad signaling, and promoting the production of the ECM [14]. Especially in the retina, TGFβ signaling demonstrates pleiotropic effects on various types of retinal cells, contributing to a wide array of functions, including the maintenance of retinal neuronal differentiation and viability, as well as the regulation of retinal vessel development and structural integrity [15]. PFD was approved in 2011 by the European Medicines Agency (EMA) and in 2014 by the U.S. Food and Drug Administration (Esbriet^®^) for the treatment of fibrotic diseases such as idiopathic pulmonary fibrosis in adults. Idiopathic pulmonary fibrosis is a chronic fibrotic and inflammatory lung disease characterized by release of pro-inflammatory cytokines, such as TNF-α and IL-1β. Although the exact mechanism of action is not fully understood, the activity of connective tissue growth factor (CTGF), platelet-derived growth factor (PDGF), α-smooth muscle actin (α-SMA), and transforming growth factor β (TGF-ß) is inhibited in the lung, as is the production of TNF-α, which plays an important role in inflammation [16,17]. It has also demonstrated interactions with a range of other molecules, including collagen I, PDGF, IL-6, IL-1β, IL-13, IL-12p40, fibronectin, HSP47, and ICAM-1, either directly or indirectly [18]. PFD also reduces fibroblast proliferation and the formation of fibrosis-associated proteins, such as collagen and fibronectin ^17,19^. In ophthalmology, animal investigations suggest that PFD may have antifibrotic therapeutic potential for proliferative vitreoretinopathy (PVR) and could offer a promising approach to treating corneal haze [20–22]. Furthermore, in another experimental mouse model focused on choroidal neovascularization (CNV), it was shown that the inhibition of TGF-β2 with PFD could offer a promising avenue for addressing choroidal neovascular fibrosis [14].

In this study, the laser-induced retinal degeneration model led to photoreceptor loss with minimal rupture of Bruch’s membrane followed by ECM formation and consequently scar formation in a focal area. After laser photocoagulation, we studied gene and protein levels of ECM proteins in the damaged retina. Furthermore, we investigated the impact of inhibiting inflammation and development of fibrosis with PFD on the formation of the ECM during retinal scar formation.

## Results

### Extracellular matrix and fibrosis development in laser-damaged retina over time

In order to quantify the laser-induced damage in the ONL, the mouse fundus was imaged using infrared (IR), OCT b-scan images were recorded and H&E overview staining was performed at various time points, which revealed an increased damage to the ONL with scarring over time (**Fig. 1A**). Live images were taken every week from 12 hours post injury to day 49 to visualize the wound healing. Quantification of the lesion area using OCT measurements of 60 lesions of 12 eyes from six mice showed a consistent reduction in lesion size until day 14. Afterwards, no significant reduction could be detected between day 14 and day 49 (**Fig. 1B**). However, a significant difference was only documented when comparing day 1 and day 5 following laser induction. Furthermore, day 5 compared with day 49 showed a substantial reduction in lesion size. However hyperintensity in OCT is stronger with time indicating fibrosis.

**Fig. 1:**
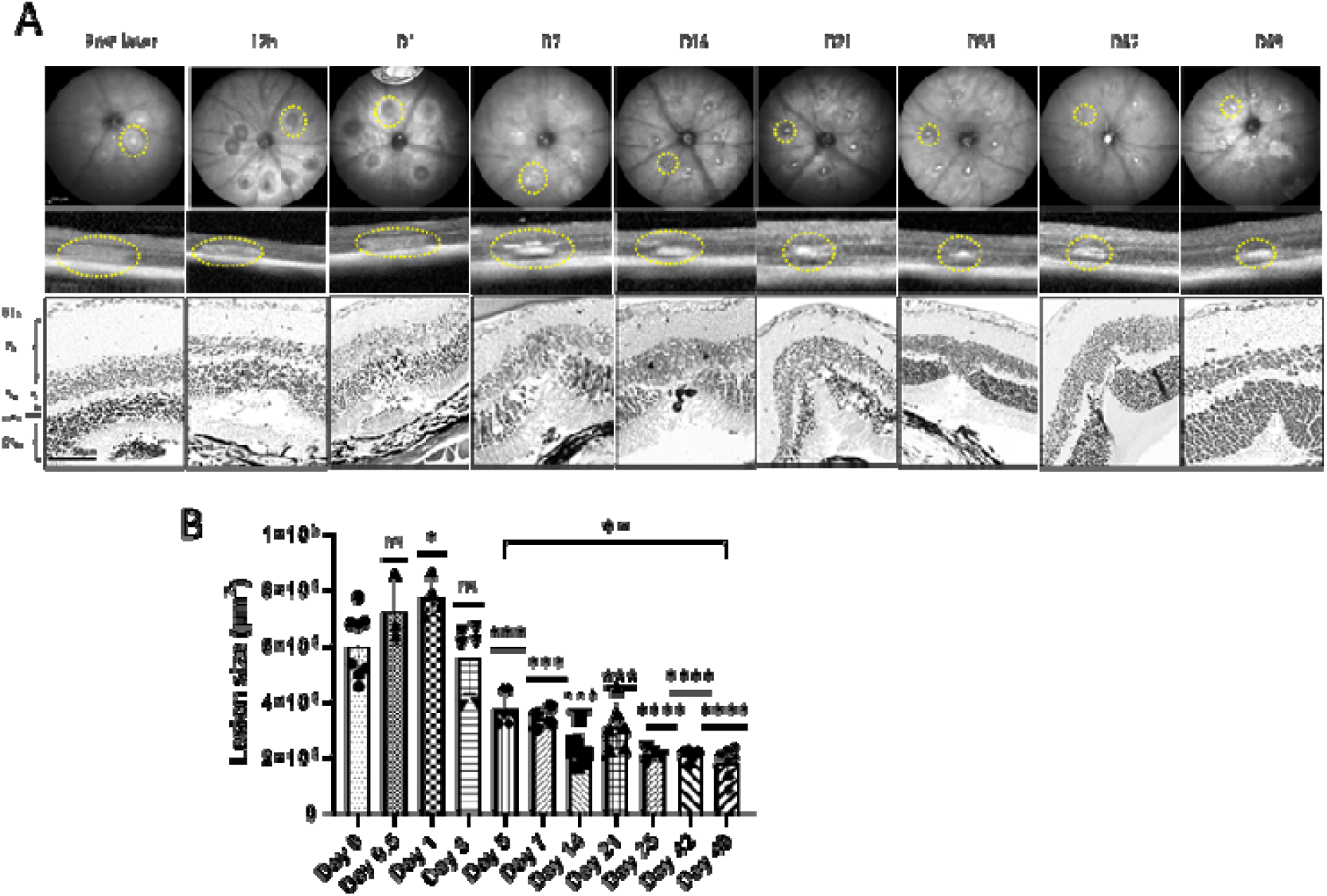
Fundus images of mice retina after photo coagulation with segmentation of OCT and representative retinal sections stained with H&E. (A) Fundus photography was taken using infrared (IR) (top) (scale bar is 200 mm) and SD-OCT (middle) at 12 hours post laser, as well as days 1, 7, 14, 21, 35, 42, and 49 post laser. H&E (bottom) shows time-dependent formation of retinal scar with photoreceptor loss (yellow circle). (B) Quantification of the lesion area on all designated time points (n = 60 lesions of 10 eyes of 6 mice per time point). The significance was analyzed with one-way ANOVA with Bonferroni multiple comparisons test for *P* = 0.05 *, *P* = 0.01 **, *P* = 0.0001 ***, *P* < 0.0001 ****

To limit new vessel growth, the laser settings were modified to perform soft tissue damage, and isolectin B4 was stained on day 7 post laser to confirm absence of CNV at the earliest detected time point (Supplementary Figure 1).

High expression of fibronectin was detectable on days 1, 7, and 14 post laser, decreased afterwards with its lowest expression on day 21 and then peaked on day 35 post laser (white outlined areas in **Fig. 2**). Moreover, fibronectin was found mainly in a scaffold in the RPE layer below the injury 12 hours post treatment. The expression of fibronectin was then found in all retinal layers at later time points from day 1 until day 49, but was detected in the RPE layer again on day 49 (**Fig. 2A**). On day 35 post laser it was not present in the entire retina anymore with the expression restricted to the damaged area only on day 49. Western Blot data also indicated a pattern fibronectin protein expression across the entire retina with slight decrease until day 21 post-damage and subsequent more obvious decrease after day 35 (**Fig. 2C,D**).

**Fig. 2:**
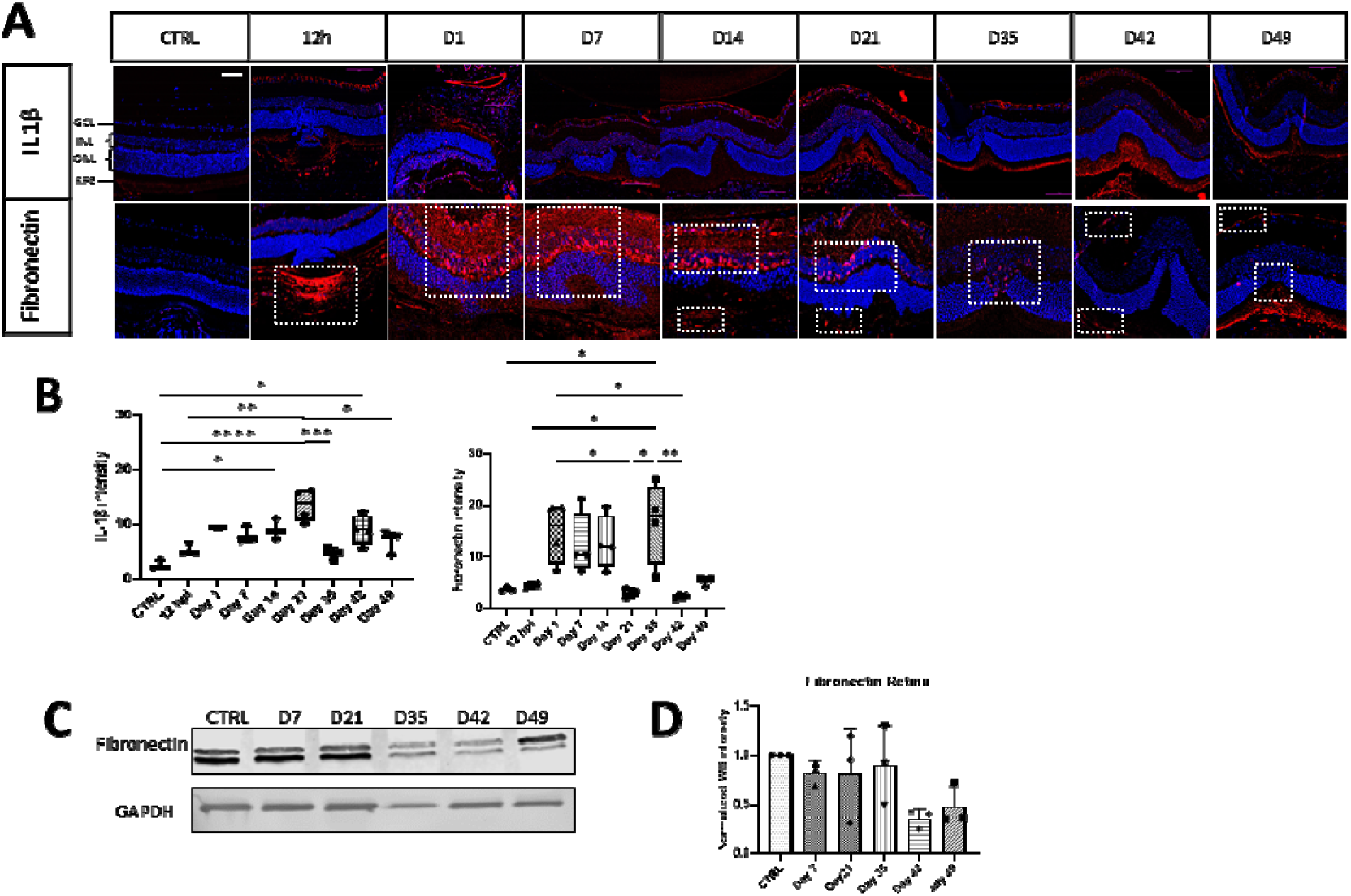
Time dependent expression of ECM marker. (A) Representative retinal section of control (CTRL) and lasered retina on selected time points stained with specific antibodies against IL-1β and fibronectin (white outlined) shown as merged images with DAPI staining. Scale bar is 50 µm. In B), fluorescence intensity was shown additional to the staining for every antibody. (C&D) Western blot analysis of fibronectin was performed. Relative protein quantification revealed a significant effect. Values are indicated as mean_J±_JSD. One-way ANOVA with Bonferroni multiple comparison test and * = *P* < 0.05, ** = *P* < 0.01, *** = *P* < 0.001, **** *P* < 0.0001, n_J=_J3/group. Scale bar: 50 µm

Additionally, the fibrotic response in the retina showed IL-1β expression over time starting from day 1 until the last time point investigated (**Fig. 2**). Thereby, active inflammation reached its maximum on day 21 with a second wave of inflammation on day 42 post laser. Interestingly, IL-1β and fibronectin increase similarly at the beginning of fibrosis (Days 1-14) but are expressed in an opposite way in the late phase (Days 21 - 49). This underscores an interplay of ECM deposition and immune response.

Immunostaining showed that the signal intensity of the activated myofibroblast marker αSMA peaked on day 21, was reduced on day 35 and slightly increased again at day 42 post laser (Fig. 3A,B). This marker is an indicator for stress fiber formation to synthesize collagens as well as other ECM components. In contrast, different collagens present in scar tissue as fiber-forming collagens 1, 3 and 5 as well as the non-fiber-forming collagen 4 increased later than αSMA detection, from 21 days until 49 days post laser (**Fig. 3A,C**). Collagen 1 reached a maximum on day 49 similar to collagen 5, whereas collagen types 3 and 4 showed their expression maximum on days 21 an 35 post laser, respectively (**Fig. 3B,D**). Apart from collagens, the multifunctional proteoglycan protein Versican demonstrated a significant increase in expression in the retina, with the highest levels observed on day 40 post damage, as detected through Western Blot analysis (**Fig. 3C,D**).

### Extracellular matrix-related genes show strong dynamics during tissue replacement

To analyze ECM-involved genes, we used the RT² Profiler™ PCR Array Mouse Extracellular Matrix & Adhesion Molecules array to profile 84 related genes (Supplementary Table 1) simultaneously for 5 selected data collection points (CTRL, day 7, day 21, day 35, and day 49). Βeta2-microglobulin (B2m) was selected as the optimal internal control for the candidate genes provided in the PCR array using the Gene Globe program offered by QIAGEN.

### General comparison

Among the 84 genes studied, the majority (70 genes) were expressed at higher levels compared to the CTRL samples on day 49 post laser, whereas 73 genes were expressed at lower levels compared to CTRL samples on day 21 post laser (**Fig. 4**). From these genes, only five (*Col4a3*, *Fbln1*, *Itgal*, *Icam1,* and *Vcan)* showed statistically significant overexpression between CTRL and day 49, the latest time point investigated (n = 3, *P* > 0.05). Additionally, *Vcan* showed a significant downregulation at day 21 (n = 3, *P* > 0.05; Supplementary Table 1). At day 7, four genes (*Cdh3, Icam1, Sell, Vcan)* showed significant different expression compared to the CTRL, while five genes (*Col3a1, Col4a1*, *Col4a3, Fbln1, Vcan*) did so on day 35 (n = 3, *P* > 0.05).

**Fig. 3:**
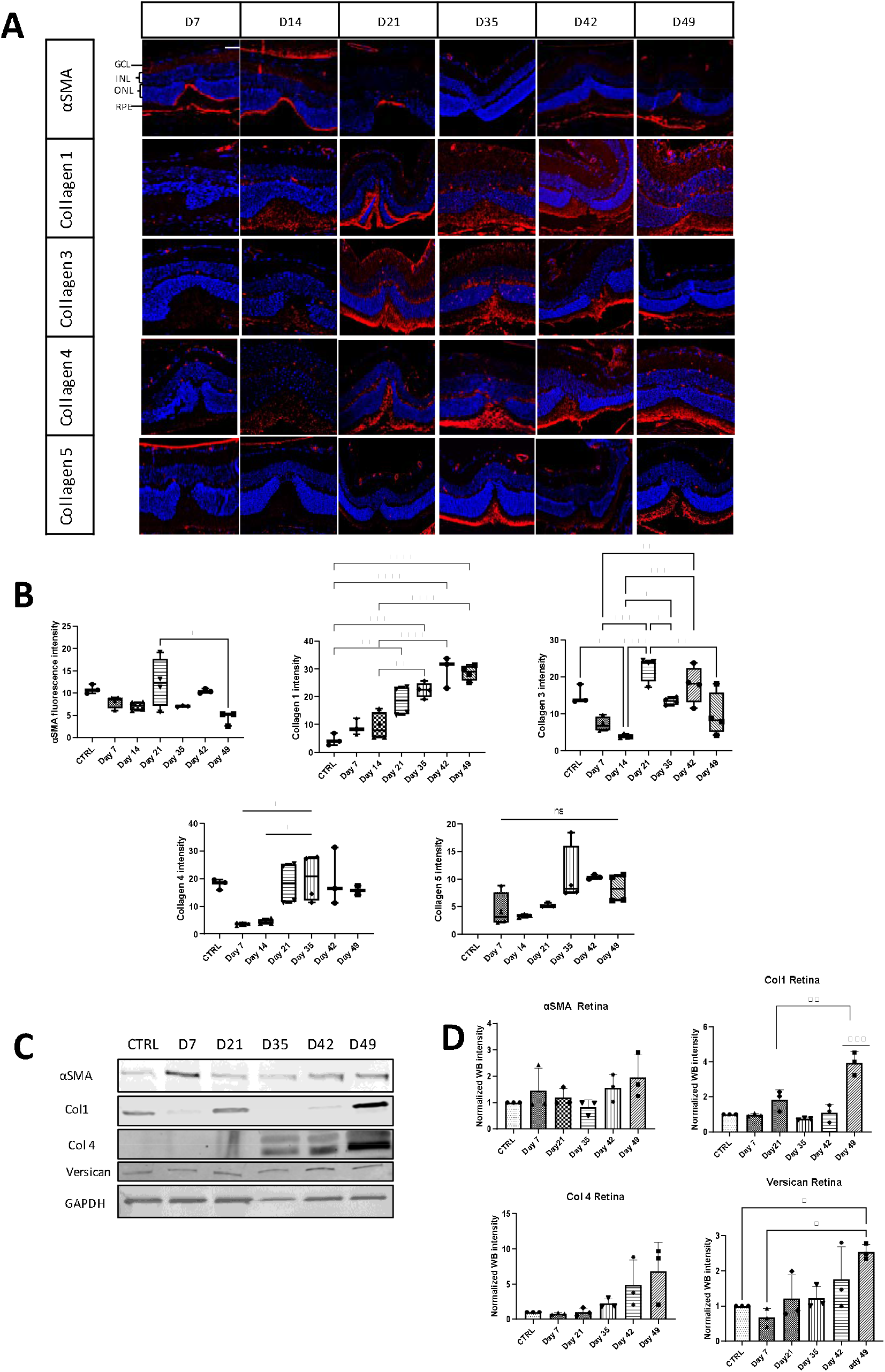
Time-dependent expression of αSMA and major component of ECM proteins in the retina of control and laser-treated mice. (A) Retinal sections were stained with specific antibodies against αSMA, collagen 1, 3, 4, and 5, and fluorescence intensity was measured for each antibody in (B). Scale bar is 50µm. (C) Western blot analysis was performed to quantify Fibronectin, αSMA, Collagen 1, Collagen 4 and Versican. (D) Statistical analysis was performed using One-way ANOVA with Bonferroni multiple comparison test, and values are expressed as mean ± SD. The study involved three mice per group, and significance levels were set at P < 0.05 *, P < 0.01 **, P < 0.001 ***, and P < 0.0001 ****.

**Fig. 4:**
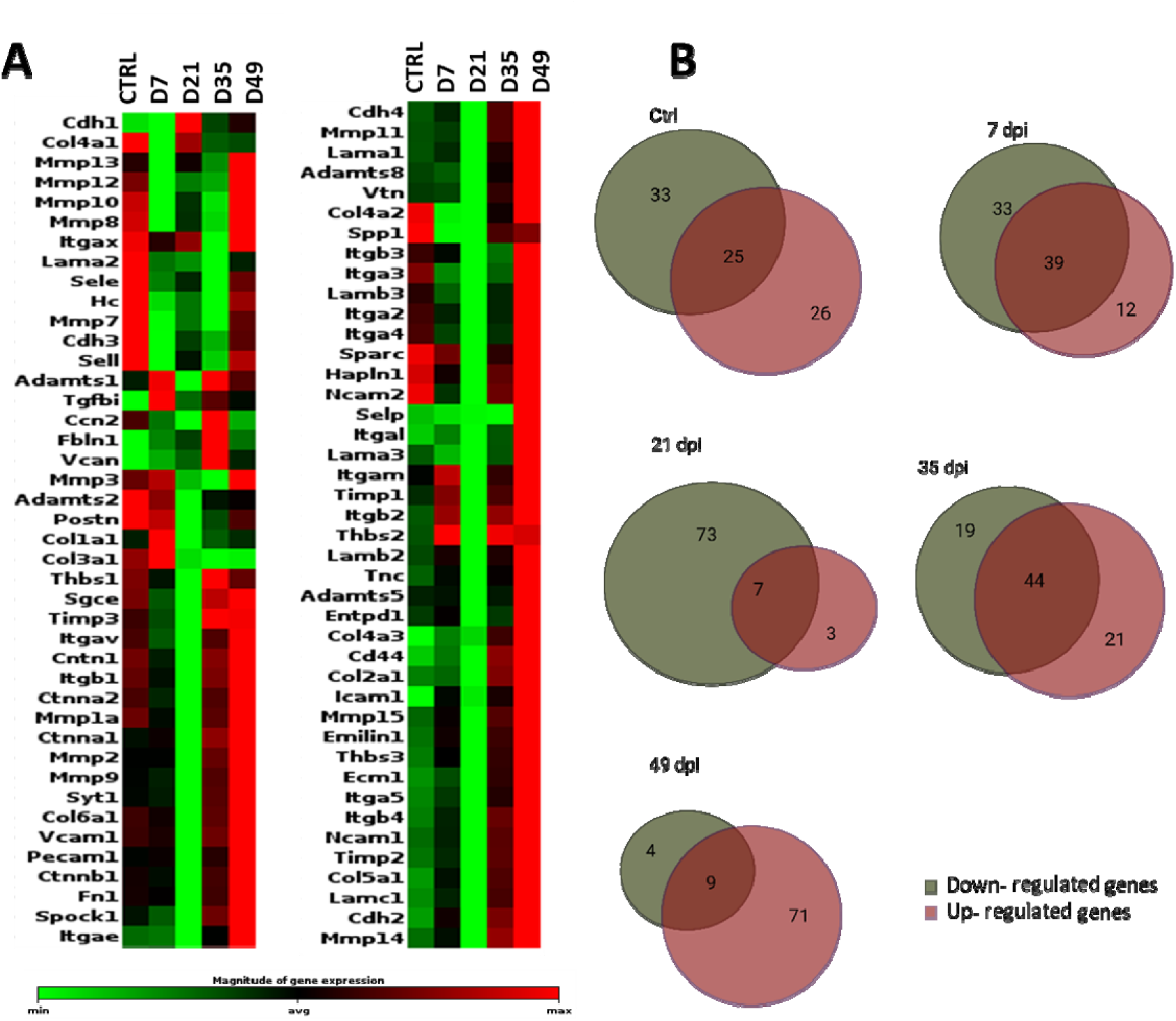
(A) Heatmap of ECM and adhesion molecule genes in the murine retina after laser injury at selected data collection points (CTRL, 7 dpi, 21 dpi, 35 dpi, 49 dpi). Differential gene expression in uninjured (CTRL) and injured retina over time is shown. RNA expression profile of grouped data from three mice (2 retinas pooled) per time point are shown as a heatmap graph. The data are displayed using a standard red-green-map, with red indicating values above the mean, black indicating the mean, and green indicating values below the mean of a row (gene) across all columns (samples). The entire dataset is hierarchically clustered in a non-supervised manner by this clustergram, which displays a heat map with dendrograms that show co-regulated genes. The marked area on the magnitude expression bar shows the defined average area for categorizing gene expression as up-regulated, downregulated, or neutral. (**B**) Venn diagrams summarize gene expression per time point grouped in up-regulated, downregulated, or neutral genes.

### Extracellular Matrix Components

Twenty-six genes included in the array (*Col1a1, Col2a1, Col3a1, Col4a1, Col4a2, Col4a3, Col5a1, Col6a1, Ecm1, Emilin1, Fbln1, Fn1, Hapln1, Lama1, Lama2, Lama3, Lamb2, Lamb3, Lamc1, Sparc, Spock1, Spp1, Syt1, Tnc, Vcan, Vtn)* were clustered as a separate group of ECM matrix proteins (**Fig. 5A,B**; Supplementary Table 1). Considering all time points, *Vcan* was significantly upregulated on days 35 and 49 (n = 3, *P* > 0.05). Furthermore, all collagen genes were significantly downregulated on day 21 (n = 3, *P* ≤ 0.05). *Vcan* and fibulin 1 (*Fbln1*) were the only two genes that were substantially upregulated at this time point (n = 3, *P* ≤ 0.05). In comparison, 18 genes showed a trend towards upregulation (*Col4a3*, *Fbln1,* and *Vcan* significantly so) on day 49 (n = 3, *P* > 0.05). At this time point, only *Col2a1* was downregulated (*P* = 0.176). On day 35, *Col4a3* and *Fbln1* were significantly upregulated (n = 3, *P* > 0.05).

**Fig. 5.**
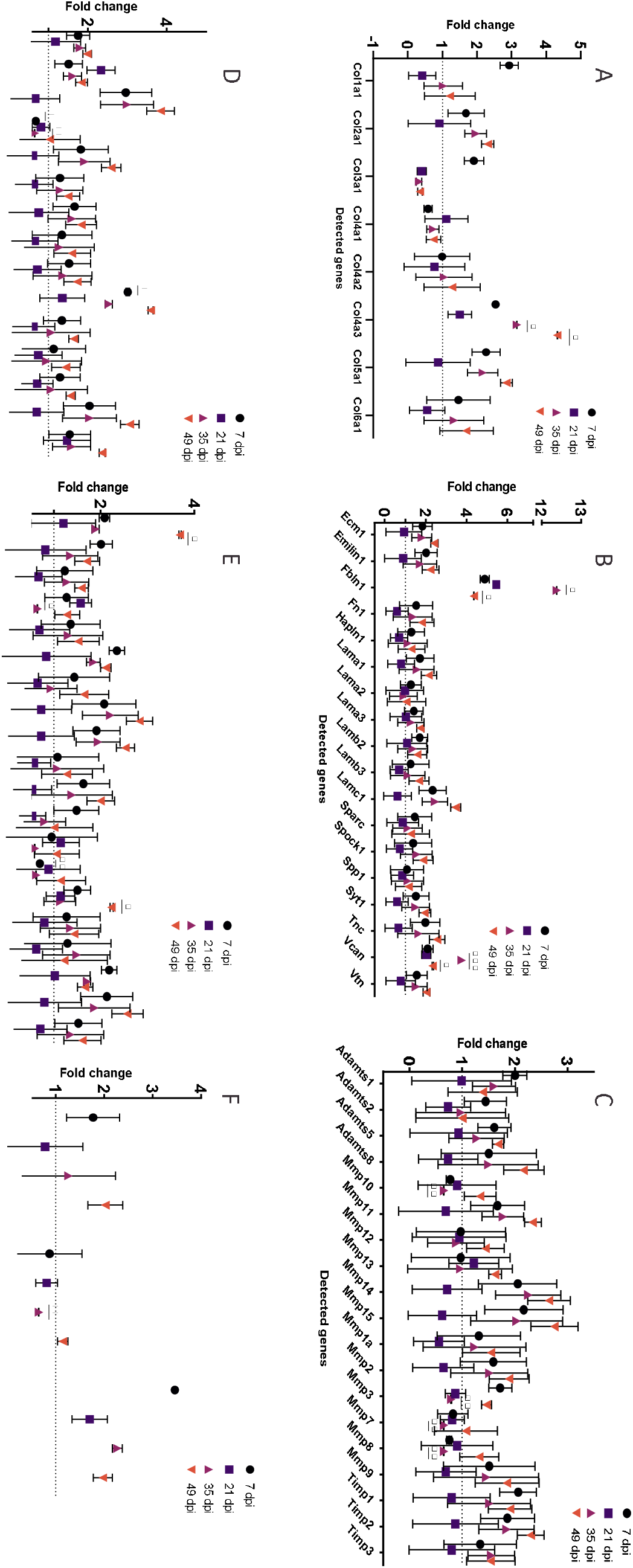

### Transmembrane Molecules and Adhesion Molecules

Another separate group of transmembrane and adhesion molecule proteins contained 35 genes. On day 7, 18 of these genes were upregulated compared to the CTRL. However, only *Icam1* was substantially upregulated (n = 3, P = 0.05; **Fig. 5D,E**; Supplementary Table 2). The only upregulation at day 21 was found with *Cdh1* (n = 3, *P* > 0.05). *Cdh2*, *Icam1*, *Itga5*, *Itgb4*, *Mmp14,* and *Mmp15* were upregulated on day 35 (n = 3, *P* > 0.05), while *Cdh3*, *Sele*, and *Sell* were significantly downregulated (n = 3, *P* > 0.05). The highest upregulation was shown on day 49 post laser; 23 of 29 genes in this subgroup were higher compared to CTRL (*Selp* and *Itgal* significantly n = 3, P = 0.03).

### Proteases and Inhibitors

Of the 19 protease and inhibitor genes investigated, only three showed significant differences compared to CTRL at any time point. However, *Timps1*, *Timps2*, and *Mmp11* were upregulated on day 7 (n = 3, *P* ≤ 0.05). Furthermore, *Mmp 10*, *Mmp3*, *Mmp7,* and *Mmp8* showed a downregulation at day 35 post laser (n = 3, *P* ≤ 0.05), while *Adamts8* and *Mmp11* were upregulated but without significance on day 49 post laser. *Mmp15* and *Mmp14* showed a trend of upregulation at all selected time points, except on day 21 post laser (**Fig. 5C**; Supplementary Table 3).

### Other ECM Proteins

Three genes included in the array (*Entpd1, Hc, Tgfbi)* did not fit in one of the previous mentioned group and were separately gathered. None showed significant differences compared to CTRL at any time point. *Tgfbi* shows a trend for upregulation at day 7 (n = 3, *P* = 0.06) and also on the other detected time points (**Fig. 5F**; Supplementary Table 4). Also *Entpd1* showed on day 7 and on day 49 post laser upregulation (n = 3, P ≤ 0.05).

### Treatment with pirfenidone alters the cell count in the outer nuclear layer and results in reduced scar formation

To determine whether PFD changed the lesion size, we quantified the volume of the laser spot with OCT measurements (b-scan, IR or AF). Comparing all collected data points of PFD-treated samples with lasered only animals (CTRL, day 21, 35 and 42), we observed reduction in lesion size (**Fig. 6A,B**). Thereby, a significant reduction comparing among all PFD samples on day 21 as well as on day 42 when comparing treated versus untreated samples. Interestingly, when PFD treatment ended on day 35 post laser, the lesion size increased again until day 42 to a comparable size as the untreated sample (**Fig. 6C**). Additional H&E was performed at the selected time points to analyze cell numbers in the GCL, INL, and ONL over time (**Fig. 6D,E**). By treating animals with PFD, a significant increase of nuclei could be only detected in the ONL at day 35.

**Fig. 6:**
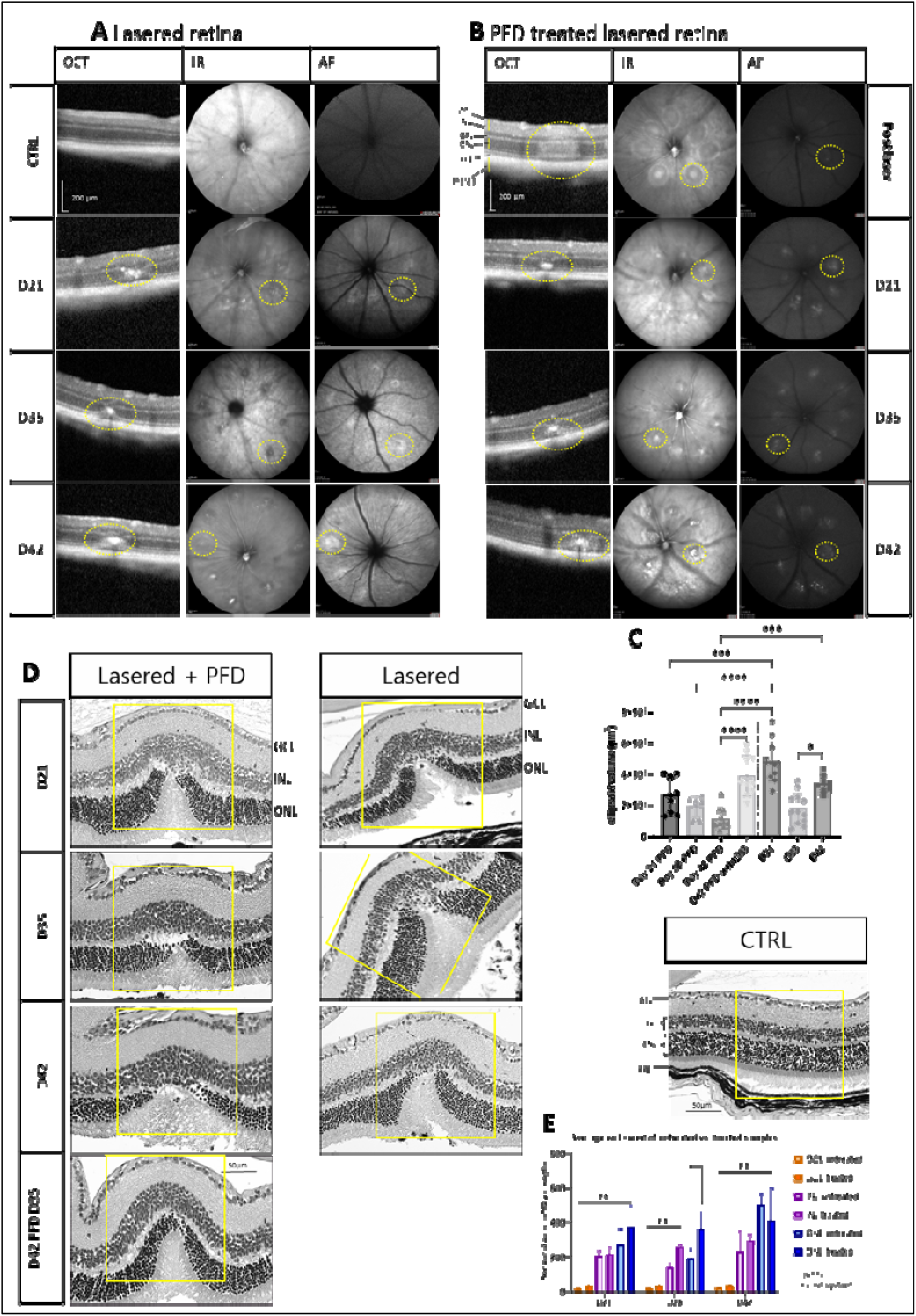
PFD treatment affects kinetic of retinal regeneration. *In vivo* imaging of lasered retina (A) and additional pirfenidone treatment from day 14 post laser (B). Fibrotic tissue was identified as hyper-reflective spot (yellow outlined). (C) The ellipsoid volume was analyzed with one-way ANOVA with Bonferroni multiple comparisons test for *P* = 0.05 *, *P* = 0.01 **, *P* = 0.0001 ***, *P* < 0.0001 ****. (D) Hematoxylin and eosin staining showed thinning of the retina. (E) Quantification of cell nuclei in ONL, INL, and GCL in lasered and additional PFD treated samples was determined using one-way ANOVA with Bonferroni multiple comparisons test for *P* = 0.05 * and n. s. (not significant). Scale bar is 50 µm.

### Pirfenidone prevents changes in collagen expression during fibrotic development

To characterize the changes in the fibrotic response after PFD treatment, we harvested retinas at days 21, 35, and 42 post laser to perform WB, rtPCR, IHC. After drug treatment, we found a reduction in collagen expression as shown in **Fig. 7**. Levels of collagen types 1, 3, and 4 were unchanged compared to non-lasered controls (**Fig. 7A,B**) and showed no increase in collagen expression as found without PFD treatment as previously depicted (**Fig. 3**). In line with these findings, quantitative Western blot analysis confirmed no significantly changed collagen protein levels in the PFD-treated samples over time. Furthermore, Western blot data showed comparatively low collagen levels as seen in IHC (**Fig. 7C**). Similar protein levels were observed for fibronectin in both Western blot and IHC at all detected time points compared to intact retina control samples. The inhibitory effect of PFD on the immune response was seen as no significant changes in IL-1β expression, neither on protein nor gene level compared to CTRL samples (**Fig. 7A,E**). This is in contrast to previously detected changes after laser treatment (**Fig. 2**). However, the activated myofibroblast marker αSMA was increased and localized in the damaged area at all detected time points until day 42 post laser. Nevertheless, neither Western blot analysis nor IHC showed significantly higher protein levels and fluorescence intensity compared to the CTRL.

**Fig. 7:**
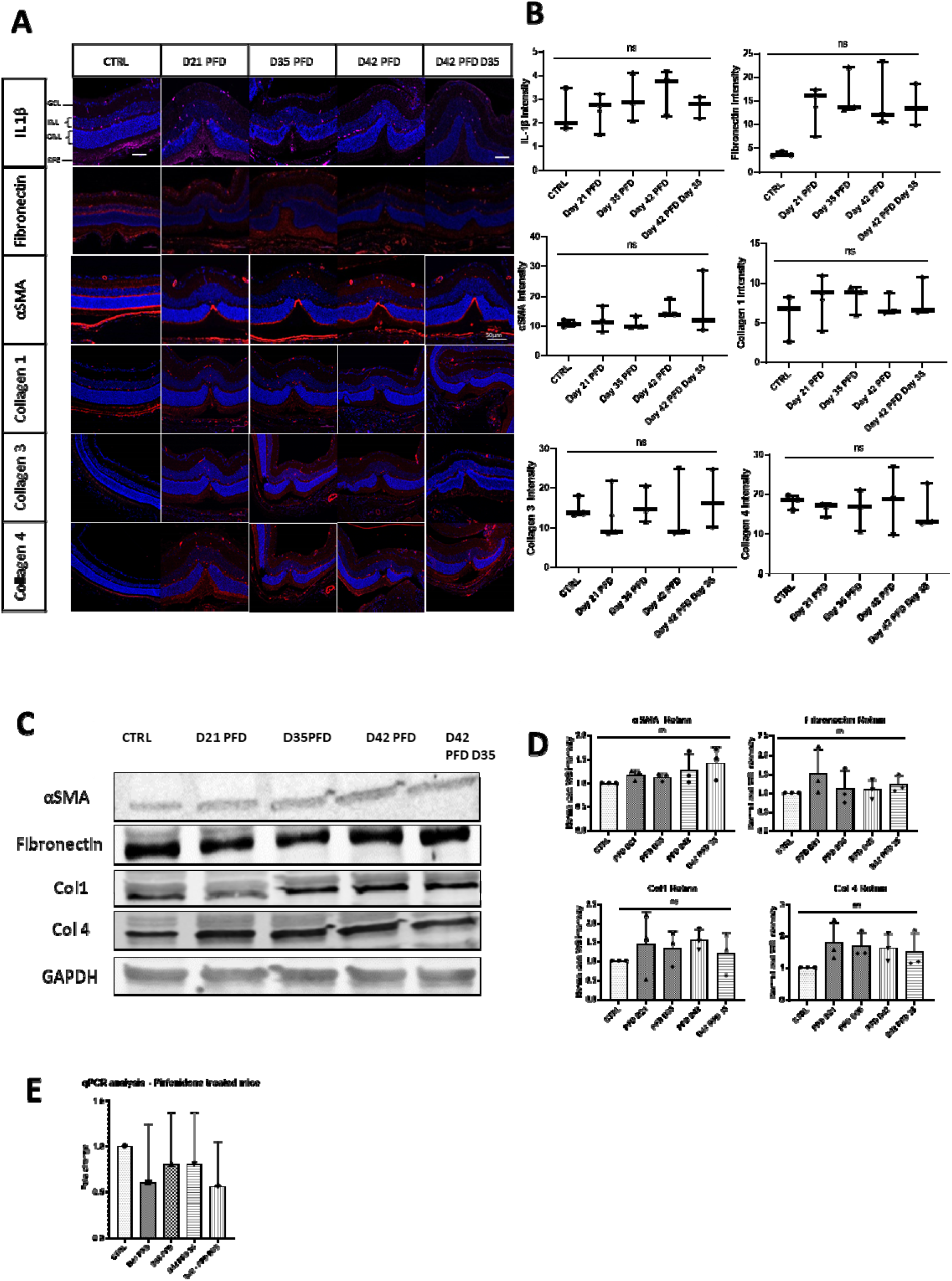
Time-dependent retinal expression of ECM markers. (A) Representative retinal section of control (CTRL) and lasered retina on selected time points stained with specific antibodies against IL-1β, fibronectin, αSMA, collagen 1, 3, 4, and 5 shown as merged images with DAPI staining. Scale bar is 50 µm. (B) Fluorescence intensity was shown in addition to staining for each antibody. (C) Western blot analysis of fibronectin, αSMA, collagen 1, and collagen 4 was performed. Relative protein quantification did not show significant changes of the investigated markers (D). (E) mRNA expression of IL-1β showed no significant changes compared to control (CTRL).

### Screening of ECM and adhesion molecules involved in the ECM formation during damage response after PFD treatment

Herein, we analyzed gain the 84 genes related to fibrotic response and extracellular matrix performance (see Table 1) to show the influence of PFD on ECM formation at the selected data points (CTRL, day 21 PFD, day 35 PFD, day 42 PFD, and D42 PFD D35).

**Table 1:** Primary antibodies used for immunostaining (IHC) and Western blot (WB) analysis.

| Antibody | Species | Type | Dilution | Company | Catalog. No. |
| --- | --- | --- | --- | --- | --- |
| αSMA | Rabbit | monoclonal | 1:200 (IHC) | Novus Biologicals | NBP2-67436 |
|  |  |  | 1:1000 (WB) |  |  |
| GS | Rabbit | monoclonal | 1:200 (IHC) | Abcam | ab16802 |
| Isolectin | *Griffonia* |  | 1:500 (IHC) | ThermoFisher | I21412 |
| GS-IB4 | *simplicifolia* |  |  | Scientific |  |
| GFAP | Mouse | monoclonal | 1:1000 (IHC)<br>1:1000 (WB) | Novus Biologicals | NBP1-05197 |
| Collagen 1 | Rabbit | polyclonal | 1:200 (IHC)<br>1:1000 (WB) | Novus Biologicals | NB600-408 |
| Collagen 3 | Rabbit | polyclonal | 1:200 (IHC) | Novus Biologicals | NB600-594 |
| Collagen 4 | Rabbit | polyclonal | 1:200 (IHC)<br>1:1000 (WB) | Novus Biologicals | PA1-28534 |
| Collagen 5 | Rabbit | polyclonal | 1:200 (IHC) | Novus Biologicals | NBP1-68938 |
| Fibronectin | Rabbit | polyclonal | 1:200 (IHC)<br>1:1000 (WB) | Novus Biologicals | NBP1-91258 |
| GAPDH | Mouse | monoclonal | 1:1000 (WB) | Novus Biologicals | NB300-221 |
| IL-1β | Rabbit | polyclonal | 1:200 (IHC) | Abcam | ab9722 |
| Versican | Rabbit | monoclonal | 1:200 (WB) | Novus Biologicals | NBP2-75706 |

### General comparisons

Among the 84 genes studied after PFD treatment, we found a balanced expression of proteins with similar amount of up- and downregulated genes. At day 21, 27 genes were downregulated and 26 genes were upregulated, whereas 28 genes were downregulated and 27 genes were upregulated at day 42 (**Fig. 8**). Of these time-listed genes, 15 showed statistically significant differences compared to CTRL at the selected time points (n = 3, *P* > 0.05). Two genes (*Col3a1* and *Col4a1*) showed a significant down-regulation and three genes (*Mmp8*, *Selp,* and *Tgfbi*) showed a significant upregulation (n = 3, *P* > 0.05) at all time points (**Fig. 9**).

**Fig. 8:**
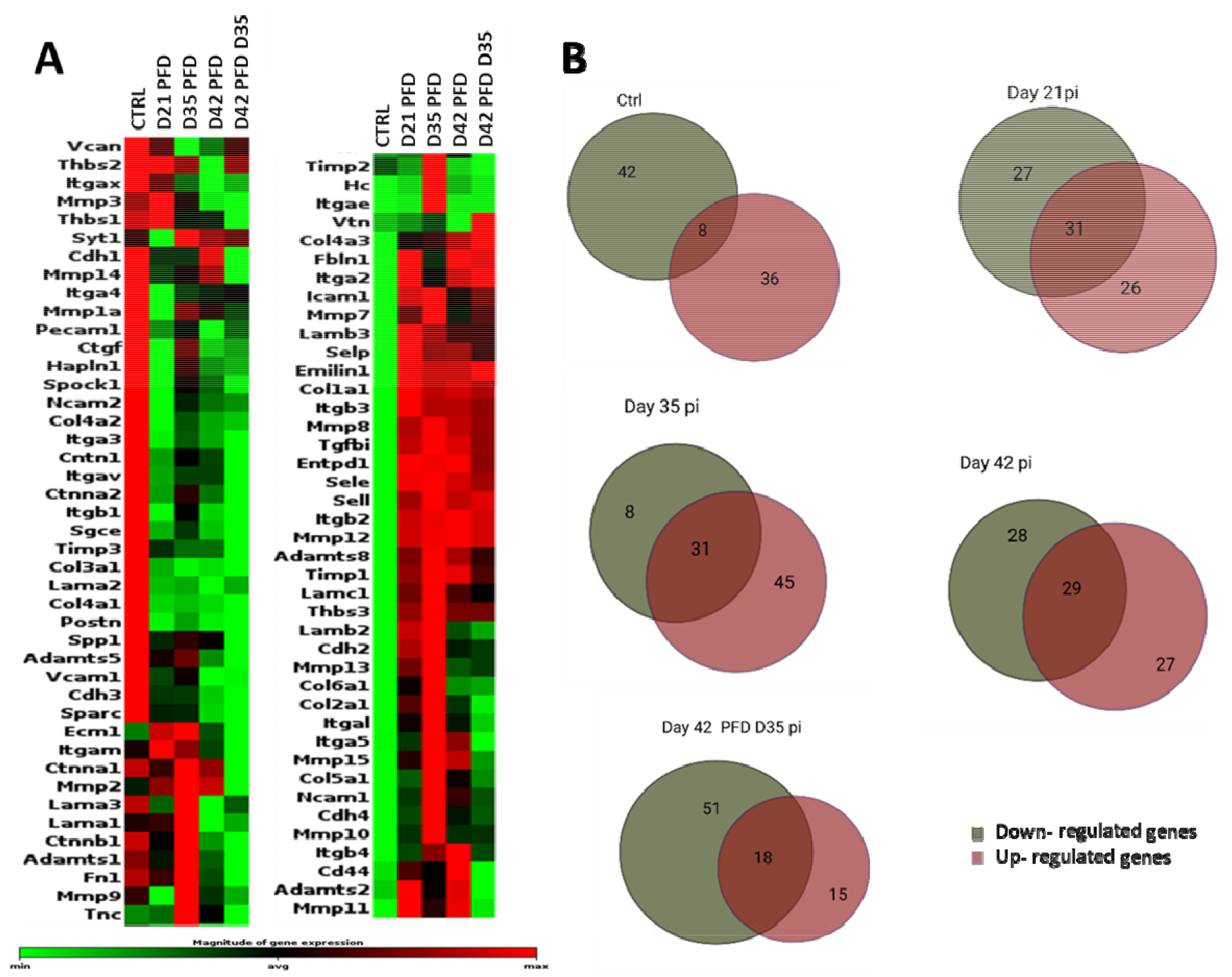
(A) Heatmap for ECM components and adhesion molecules in murine retina after laser injury in selected time points (Day 21 PFD, Day 35 PFD, Day 42 PFD, and Day 42 PFD D 35 post injury). Differential gene expression in normal (CTRL) and injured retina over time is shown. Grouped data from three mice retinas (2 retinas grouped) per time point are shown. The data are displayed using a standard red-green-map, with red indicating values above the mean, black indicating the mean, and green indicating values below the mean of a row (gene) across all columns (samples). Marked area on the magnitude expression bar shows defined average area for grouping gene expression into up- regulated, downregulated, or neutral. (B) Venn diagrams summarize gene expression per time point grouped in up- regulated, downregulated, or neutral genes.

**Fig. 9.**
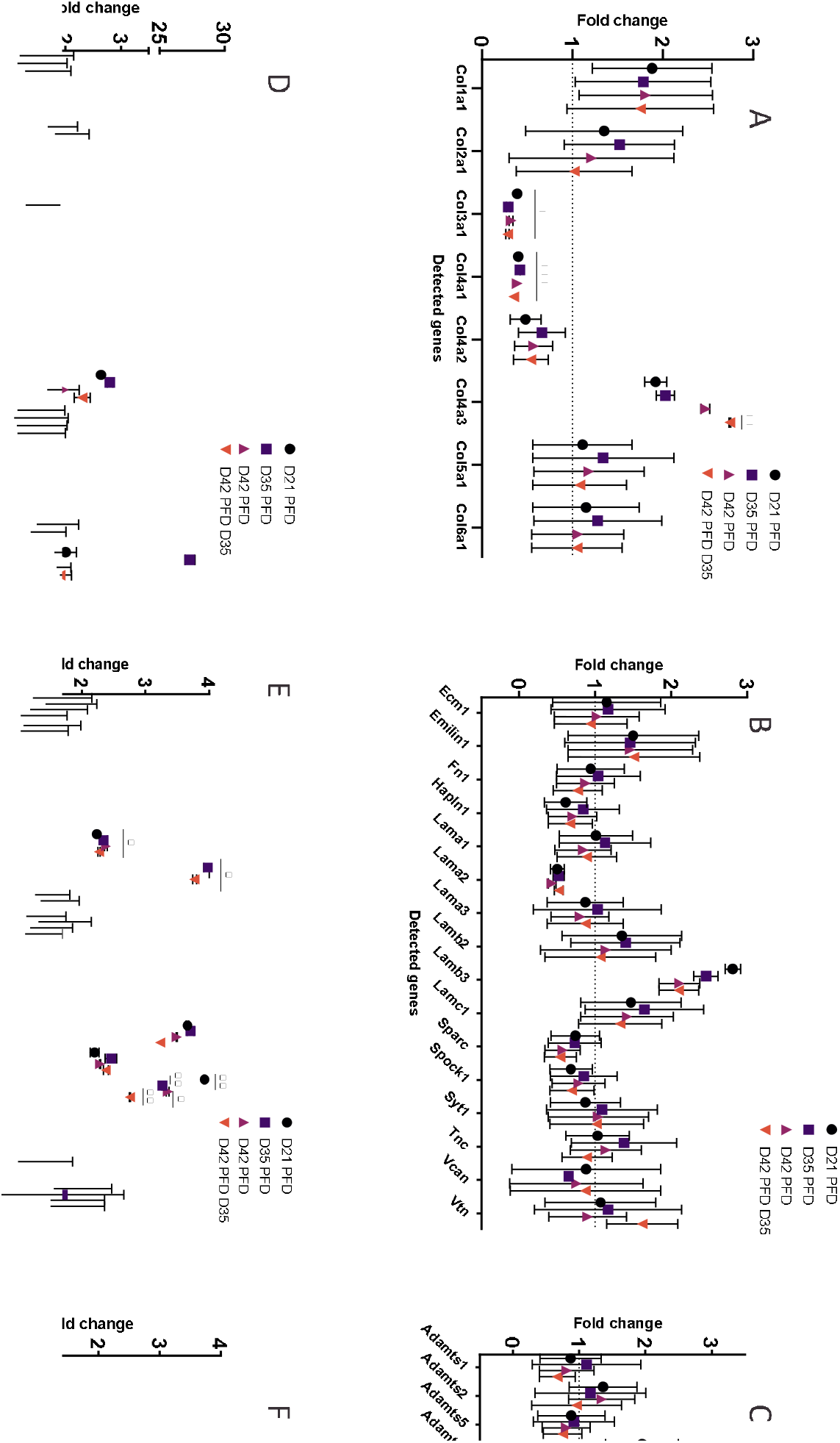

### Extracellular Matrix Components

*Col3a1* and *Col4a1* were significantly downregulated compared to the CTRL (n = 3, *P* > 0.05) as was *Lama2*, but without significance (n = 3, *P* > 0.05) at all time points (**Fig. 9A,B**; Supplementary Table 1). The analysis showed significant upregulation of *Col4a2* and *Lama2* on day 21. Moreover, *Col1a1*, *Lamb3,* and *Col4a3* showed an upregulation at all selected time points (*P* ≤ 0.05). Both PFD treated animal groups until day 42 were analyzed and depicted an upregulation for *Col1a1* without significance. *Emilin1* shows a trend and was slightly increased at all detected time points (*P* = 0.8).

### Transmembrane Molecules and Adhesion Molecules

From 34 genes, *Itgb2*, *Itgb3*, *Sele*, *Sell,* and *Selp* were upregulated at all time points. *Sele*, *Itgb2,* and *Itgb3* showed a trend and were increased on day 21, day 42, and day 42 PFD day 35. *Sell* was increased for both samples of day 42 (n = 3, *P* ≤ 0.05), while *Selp* showed significant upregulation at all time points (*P* = 0.02; **Fig. 9 E,F**; Supplementary Table 6).

### Proteases and Inhibitors

*Mmp8* was significantly upregulated at all time points, as was *Mmp12*, but only significantly at day 21 and D42 PFD D35 (**Fig. 9 C**; Supplementary Table 7). Furthermore, the *Timp* genes showed a non significant upregulation at all investigated time points (n = 3, *P* > 0.05).

### Other ECM Proteins

Only *Tgfbi* was significantly upregulated at all detected time points compared to the CTRL (n = 3, P > 0.05; **Fig. 9F**; Supplementary Table 8).

## Discussion

Retinal wound repair is a complex process, which involves interconnected biological mechanisms, including inflammation, cell proliferation, and tissue remodeling. In general, four overlapping phases in the process of general wound healing have been defined. The first phase is characterized by hemostasis, during which the bleeding is stopped. The second phase is an immediate inflammatory response that involves the infiltration of leukocytes, which release cytokines with antimicrobial properties. These cytokines activate the third phase, the proliferative phase, in which amongst others new ECM is generated. Finally, in the fourth phase, the wound contracts over a period of weeks to months as the ECM is restructured [23]. These specific changes to the ECM during remodeling of the retina after the loss of photoreceptors are depended on the complex of several cellular and molecular components, which are not fully understood. Thereby, the function of the ECM is not only structural support, it regulates also fluid and solute passage from the choroid to the retina. In this study, the expression of key ECM components after laser-induced damage in adult mouse retina was investigated by inducing photoreceptor degeneration and retinal thinning that occurs in many retinal diseases. This time-dependent model enabled spatial observation of internal processes during retinal scarring caused by photoreceptor loss. The herein employed laser model differs from a recently published CNV model, which leads to subretinal fibrosis [24]. In comparison, the subretinal fibrosis model published by Zandi at al., we showed absence of neovascularization at all time points post laser by negative isolectin B4 staining. In contrast, therein an increase in CNV until day 14 and a regression of CNV with an increase of subretinal fibrosis after day 21 post injury by initially rupturing Bruch’s membrane and inducing a characteristic bubble, was displayed.

Current research has shown that several types of retinal cells, including RPE cells, endothelial cells, epithelial cells, Müller cells, and macrophages, have the ability to transform into myofibroblasts. These myofibroblasts are the primary cell type responsible for triggering fibrosis and producing ECM components [1,25,26]. They also express α-SMA, a filamentous contractile protein, during activation state and their persistent stimulation is associated with excessive ECM deposition leading to organ dysfunction [27]. Our data show that α-SMA expression displays biphasic kinetics after laser-induced retinal damage. However, as shown in the unlasered control, α-SMA is also present in the choroidal space under normal physiological conditions, where it plays a pivotal role in regulating vascular diameter, vascular tone, and retinal blood flow [28].

One primary response of wound healing is acute inflammation through recruitment of immune cells and fibroblasts to the site of tissue damage. However, effective wound repair necessitates the resolution of inflammation, while an excessive inflammatory response can lead to the formation of retinal scars [29]. In our model, we showed the development of three stages of cytokine secretion indicating several waves of inflammation. Firstly, an initial IL-1β production within 24 h post injury. Second, an intermediate phase with no difference compered to baseline amounts. Finally, a repeated biphasic increase on days 21 and 42. Tissue with regenerative potential resolves the inflammatory response within a few days with removal of myofibroblasts and inflammatory macrophages, so that the scar gradually breaks down within several weeks [23]. In wet AMD, the connection between inflammation and abnormal blood vessel growth is well known^1^. In dry AMD, the role of inflammation caused by drusen formation due to accumulation of toxic debris, such as lipofuscin, has been found to form a chronic and toxic milieu for RPE cells, thus creating a pro-inflammatory microenvironment [30]. As we did not induce CNV lesions that actively destabilize the Bruch’s membrane and had no drusen formation as inflammatory mediators in our mouse model, our data indicate repeated release of IL-1β in the damaged area. This might lead to the destabilizing of the BRB and leukocyte infiltration. It is also known that macroglia cells show upregulation of the pro-inflammatory cytokine S100β under stress conditions and microglia are initiators of chronic scarring [31]. However, the observed three-phasic inflammation signal influences the fibrotic response towards chronic scar formation and hindering therewith any repair process. As one of the earliest ECM matrix proteins, fibronectin appears 10 min post injury and was recently described as an inflammatory mediator [23,32]. Fibronectin expression in the RPE layer peaks at early and late stages of wound repair in the injured area. We therefore hypothesize that inflammatory mediators are active in the RPE layer as well as in retinal cells so that the complex architecture of the ECM is continuously synthesized.

Since collagens are the key components of a healing wound and are the most prominent proteins of the ECM scaffold, we investigated the appearance of collagen types 1, 3, 4, and 5 via immunochemistry. While collagen types 1, 3, and 5 belong to the group responsible for fiber formation with long ropelike structures and assemble into polymers; collagen type 4 stands out as the primary constituent of the basement membrane, possesses the ability to organize into sheet-like networks within this structure [33]. Increased expression was detectable for collagen 1 from day 14, for collagen 3 from day 21, and for collagens 4 and 5 from day 35 on. The expression of all analyzed collages was found until day 49 post laser.

Similar to the observed inflammation marker IL1β, collagens showed a wavelike expression. Compared to findings in other tissues, including skin, heart and liver, our results diverge from reported observations regarding a shift in the collagen 1 to collagen 3 ratio. These changes typically manifest as an increased concentration of collagen 3 during the healing phase, followed by a subsequent rise in collagen 1 content in the healed tissue [34]. On the other hand, our data indicate a substantial presence of collagen types 1 and 3 persisting until the last recorded time point on day 49. This suggests a notable constraint on wound healing, with no discernible indications of regenerative processes. The gene expression levels showed significant increase only for *Col4a3*, even with the majority of collagens upregulated at day 7 and day 35 post laser. This result fits the characteristics of the retina as collagen 4-rich tissue, which is mainly found in the Bruch’s membrane and the inner limiting membrane and is critical for neuronal survival and angiogenesis [35]. On the other hand its abnormal deposition is seen in fibrotic lesions of certain organs including kidney and lung [36,37].

Herein, we provide an overview of the complex ECM formation in our injury model, and show that the most robust alterations in gene expression were detected for *Col4*α*3*, fibulin1 (*Fbln1*), cadherin 3 (*Cdh3*), intracellular adhesion molecule-1 (*Icam1*), integrin alpha L (*Itgal*), selectin E (*Sele*), transforming growth factor beta-induced (*Tgfbi*), and versican (*Vcan*). Except *Cdh3* and *Sele*, all genes were significantly upregulated, mainly at day 35 and day 49 post laser in the late stage of retinal fibrosis. Most of these genes indicate a causal role for leucocyte infiltration during the regenerative phase, which might inhibit remodeling. The observed upregulation of *Fibulin 1* in our wound healing model corresponds to a study on patients with pulmonary fibrosis. This study identified the bioactive region/s of fibulin-1C responsible for promoting fibrosis, indicating potential shared mechanisms in fibrotic processes [38]. A binding partner of fibulin 1 is versican, a member of the hyalectin family of ECM components. In the retina, similar to the brain, ECM is formed by proteoglycan proteins, such as versican (Vcan), to implement the three-dimensional network of extracellular macromolecules. In most diseases, including lung disease and several different cancers, these proteoglycans increase dramatically during inflammation [39]. Our data concurred as we show *Vcan* upregulation in the late stage of tissue response. The literature has indicated that neurons and glial cells might synthesize proteoglycans and glycoproteins together with fibrous proteins, such as collagens and fibronectin [40]. Thereby, proteoglycan as well as P- and L-selectins increase dramatically during inflammation and interacts with receptors that are found on the surface of immune cells, [41–43]. According to existing literature, the ECM exhibits an organization into cable-like structures involving versican and other proteins, including hyaluronan, which bind to leukocytes. Further investigations are warranted to elucidate the specific contributions of these complex components to the phenotype of leukocytes.

Furthermore, the upregulation of the protein Icam 1 is particularly essential for the migration of leukocytes and has been described in several ocular diseases, such as diabetic retinopathy and CNV [44]. In other organs, such as liver, Icam 1 has been described as an inflammatory and fibrotic marker [45]. We could detect two peaks for *Icam 1* expression at the gene level, namely on day 7 and day 49 post laser. These findings could imply not only leukocytes entering the retina but also their role in fibrosis. Another marker, *Itgal*, was significantly upregulated on day 49 post laser. This gene encodes integrin components, including lymphocyte function-associated antigen 1 (LFA-1), which are expressed on lymphocytes to participate in immune and inflammatory responses [25]. In contrast, *Cdh3* (also known as P-cadherin) was significantly downregulated on day 7 and day 35 post laser. Previous publications have shown that Cdh3 is the dominant cadherin in mature RPE cells [46]. It plays a critical role in maintaining the structural integrity of epithelial tissues, regulating processes involved in embryonic development, and maintaining adult tissue architecture and cell differentiation. The downregulation of *Cdh3* suggests a decreased density of RPE after laser damage. Thus, the two-wave dynamic of *Cdh3* downregulation might indicate a destabilized RPE layers, leading to higher permeability with migration of peripheral immune cells. Chd3 might also influence stem cell activity. Earlier studies have reported a significant loss of epithelial progenitor cells and downregulation of putative stem cell markers in organ-cultured limbal tissue [47].

To modulate the laser-induced damage including fibrosis and ensuing immune response, we used PFD, which is not only anti-inflammatory but has anti-fibrotic properties also. PFD inhibits fibroblast proliferation and the production of fibrosis-associated proteins and cytokines that increase the biosynthesis of ECM proteins, such as collagen and fibronectin. Such cytokine secretion is mediated by TGF-β or PDGF pathways [48]. Despite demonstrating a reduced expression of collagen 1, 3, 4 and 5 and IL-1β after PFD treatment, we were not able to detect long-term effects on the scar tissue. This finding suggests a yet undefined time window in which inflammation must be prevented or halted. Otherwise, even after inhibiting inflammation and reducing fibrotic signals, fibrosis inevitably occurs [49]. Day 14 post laser is likely already too late as the starting point for inhibition of fibrosis development and modulation. Alternatively, additional pro-inflammatory cytokines are produced or inflammation is not downregulated in general. Tissue with limited regenerative capacity, such as the retina, has a high risk for repetitive fibrotic responses or chronic developments with aggregation of ECM, myofibroblasts, and macrophages [50]. Examining the expression of involved genes, we found a shifting of gene expression toward 12 pro-fibrotic genes after PFD treatment compared to non-treated samples. Thereby, *Col3*α*1* and *Col4*α*1* were downregulated, whereas *Itgb2* and *Itgb3* were upregulated.

The selectin family, composed of L-selectin (SELL), E-selectin (SELE), and P-selectin (SELP), plays a significant role in mediating cell adhesion. Most leukocytes express L-selectin, while activated endothelial cells express E-selectin and P-selectin [51,52]. All three family members were significantly upregulated after PFD treatment. Based on our hypothesis, the drug treatment might activate endothelial cells in the surrounding blood vessels, which facilitate leukocyte infiltration by assisting in their adhesion and subsequent transportation into the retinal layers, thereby influencing remodeling processes.

The expression of transforming growth factor beta induced (*Tgfbi*), which is regulated by TGF-β signaling, was significantly upregulated at all detected time points. The ability of Tgfbi to bind to ECM proteins, such as fibronectin, laminin, vitronectin, and collagens, is likely due to fasciclin 1 (FAS1) [53].

Matrix metalloproteinases (MMPs) are a class of enzymes responsible for breaking down the ECM and regulating ECM turnover and homeostasis. Tissue inhibitors of metalloproteinases (TIMPs) are specific inhibitors of MMPs that bind to active MMPs and inhibit the process of ECM degradation. Previous research has shown that excessive expression of TIMP-1 may hinder wound healing by impeding epithelial cell migration, a critical aspect of re-epithelialization. Although we observed an upregulation of *Timp1* levels at all selected time points after PDF treatment, this upregulation was not significant.

In conclusion, inflammation peaks on day 1, day 21 and day 49 in the used laser-based retina degeneration model. This inflammation hindered remodeling as fibrosis continued over 49 days post laser treatment. Our findings suggested that *Cdh3* downregulation in the late stage might explain the absence of stem cell activation or proliferative capacity in the mouse retina.

However, the study had some limitations as the focus lay mainly on the mRNA expression in lasered retina samples. The samples showed a limited amount of laser damage that might explain the low significance for most of the detected genes. Additionally, we did not focus on changes in the surrounding tissues (e.g., choroid) during scarring, which may also be part of the interaction between ECM / adhesion molecules and, especially, the involved cells. Therefore, further studies are needed to validate our findings and conjectures *in vivo*.

### Experimental procedures

#### Laser-induced retinal damage

Adult C57BL/6J mice (6–12 weeks; Charles River Germany, Sulzfeld, Germany) were kept under standard conditions in individually ventilate cages (IVC) in a temperature-controlled animal facility with a 12-hour light/dark cycle. They were fed with standard laboratory chow and water *ad libitum*. All experimental procedures were approved by the governmental authorities of the Canton of Bern (BE 146/2020).

The mice were anesthetized with 1 mg/kg medetomidine (Domitor, 1 mg/mL; Provet AG, Lyssach, Switzerland) and 80 mg/kg ketamine (Ketalar, 50 mg/mL; Parke-Davis, Zurich, Switzerland). The pupils were dilated with 2.5% phenylephrine and 0.5% tropicamide (ISPI, Bern, Switzerland). Hydroxypropyl methylcellulose (Methocel 2%; OmniVision AG, Neuhausen, Switzerland) was applied to keep the eyes hydrated. To induce retinal damage in the ONL without Bruch’s membrane rupture, the eyes were lasered using a 300 μm spot size, 60 ms duration, and 60 mV pulses of an 532-nm laser photocoagulation (Visulas 532s laser workstation with slit lamp; Carl Zeiss Meditec AG, Jena, Germany). Eight laser spots were centered with 2–3 disk diameters from the optic nerve by using a coverslip to allow viewing of the posterior pole of the eye. Lesions in which bubbles, a criteria for Bruch’s membrane rupture, were identified during the laser process were excluded from the study.

#### Pharmacological treatment

To inhibit the inflammatory response via the TGFβ pathway, 10 mg/kg pirfenidone (PFD; Catalog No. S2907, Selleck Chemicals, Houston, TX, USA) was administered via drinking water. To protect the drug stability, the water bottles were covered from light. The control animals received an equal amount of water under the same conditions. For monitoring, the animals were weighed three times per week. Water consumption was measured daily and the water was thereby exchanged. The PFD treatments started on day 14 post laser injury until samples were collected (day 21, day 35, and day 42 post laser). Additionally, to investigate the influence of retinal recovery, we terminated the PFD treatment on day 35 post laser injury and harvested the retina 7 days later on day 42 post-laser. This sequence is defined as “D42 PFD D35” throughout the manuscript.

#### Histology

Paraffin-embedded sections (5 μm) of mouse eyes were deparaffinized and rehydrated through a graded series of xylol and alcohol. The sections were stained for 5 min with hematoxylin (Sigma, St. Louis, MO, USA) and 1 min with eosin (Roth, Karlsruhe, Germany) and then mounted with Eukitt® (Medite Service AG, Dietikon, Switzerland).

#### Staining

For immunostaining (IHC), deparaffinized samples were blocked at room temperature in blocking solution containing 3% normal goat serum (Agilent Technologies, Santa Clara, CA, USA), 0.5% casein (Sigma), and 0.05% Triton X-100 (Sigma) in Tris buffered saline (TBS) for 30 min. Primary antibodies (Abs) were added and incubated at 4°C overnight; these were Abs against gliosis-specific markers or ECM-relevant proteins, as summarized in Table 1. After incubation, sections were washed 20 min and stained at room temperature with the respective secondary Abs (Alexa Fluor® 488 & 594 Abcam, 1:500) and 4’,6-Diamidino-2-phenylindole (DAPI; Sigma) to counterstain nuclei for identification for 3 h. Images were acquired using a Nikon Eclipse 80i microscope (Nikon, Tokyo, Japan) at 20x magnification and processed using ImageJ 1.51n (Fiji-win64, NIH, Bethesda, MD, USA).

To determine the number of cells in the ganglion cell layer (GCL), the inner nuclear layer (INL), and the ONL, manual counting was performed within the lasered area in the hematoxylin and eosin (H&E) images. The size of the counted area corresponded to a retinal section 100 µm in length.

#### Western blot analysis

To obtain tissue, the eyes were enucleated immediately after optical coherence tomography (OCT) imaging. The retinas were microsurgically isolated from the choroid-retinal pigment epithelium (RPE) complex and two were combined in 100 μl of lysis buffer (**Table 2**). There, they were supplemented with a protease inhibitor cocktail (cOmplete™ Mini; Merck, Darmstadt, Germany) and phosphatase inhibitors cocktail (P2850; Sigma) and homogenized (Precellys® 24 tissue homogenizer; Bertin Technologies S.A.S, Montigny-le-Bretonneux, France). The lysate was centrifuged (12,000 rpm, 15 min, 4°C), and the supernatant was collected. Each sample containing 30 µg of total protein, as quantified by using the Bradford assay (ThermoFisher Scientific, Reinach, Switzerland), was separated by SDS-PAGE and electroblotted onto nitrocellulose membranes (Trans-Blot® Turbo™ Transfer System; Bio-Rad Laboratories, Cressier, Switzerland). To block nonspecific binding, the membranes were washed with milk Intercept™ (TBS) Blocking Buffer (LI-COR, Lincoln NE, USA) and subsequently incubated at 4°C overnight, followed by incubation for 3 h with a donkey anti-rabbit IgG polyclonal antibody (IRDye® 800CW) or donkey anti-mouse IgG polyclonal antibody (IRDye® 680RD), respectively. The signals were visualized by infrared laser (Li-Core Odyssey; LI-COR). Quantification was performed using Fiji-win64 (Image J; NIH).

**Table 2:** Lysis buffer ingredients.

| Ingredients | Concentration |
| --- | --- |
| Tris-HCl (pH 7.5) | 20 mM |
| NaCl | 150 mM |
| EDTA | 5 mM |
| Na-Pyrophosphate | 5 mM |
| NaH<sub>2</sub>PO<sub>4</sub> (pH7.6) | 20 mM |
| Na-β-glycerophosphate | 3 mM |
| NaF | 10 mM |

#### RT-qPCR

Total RNA from the retinas including negative controls (uninjured retina without drug treatment from age-matched siblings) was extracted at different time points (days 21, 35, and 42 post laser) after PFD treatment using the TRIzol™ (Thermo Fisher Scientific) according to the manufacturer’s instructions. Three independent samples from two pooled retinas were used for each condition. The cDNA was reverse-transcripted by using iScript cDNA Synthesis Kit (Bio-Rad) according to the manufacturer’s instructions and quantified using a NANODROP 1000 spectrophotometer (Thermo Fisher Scientific). The primer sets are summarized in **Table 3**. RT-PCR was performed in a 20 µl reaction containing cDNA specific primers with iTaq^TM^ Universal SYBR Green Supermix (Bio-Rad) using the CFX Connect™ Real-Time PCR Detection System (Bio-Rad). The thermocycling conditions were as follows: 98°C for 3 min, followed by 40 cycles at 98°C for 15 sec, 58°C for 15 sec, and 72°C for 15 sec. The relative abundance of transcripts was normalized according to the relative abundance of the reference gene glycerinaldehyd-3-phosphat-dehydrogenase (GAPDH). Gene expression was measured by the change in threshold (ΔΔCT). Expression data were presented as mean=±=SD calculated against the negative control samples, and expression in the control samples was set to 1.

**Table 3:**
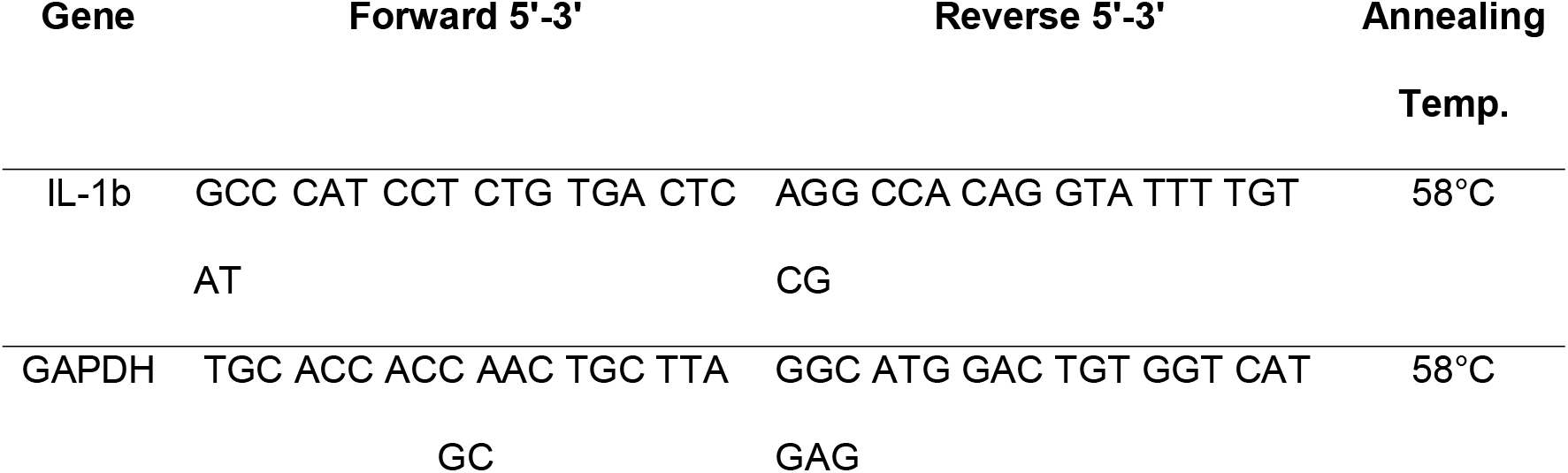
Primer Pairs for RT-qPCR.

**Table 4:**
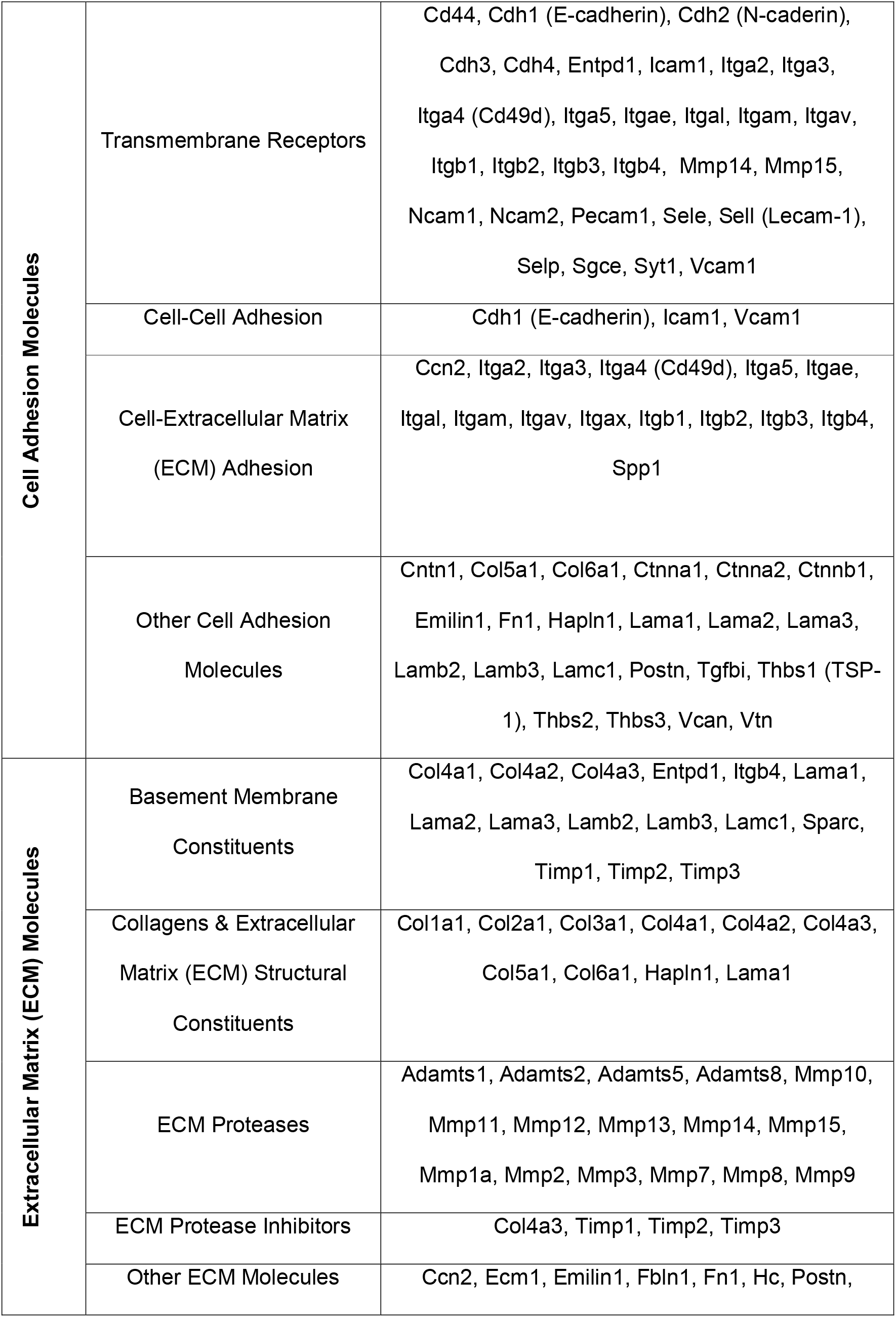
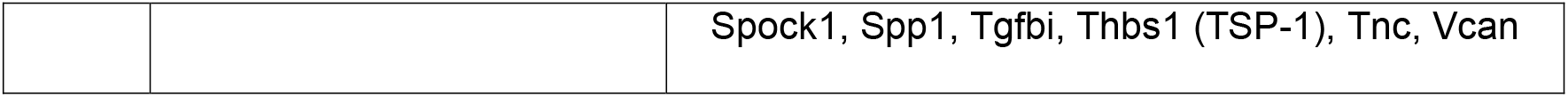
Summary of genes detected in the RT^2^ ECM and adhesion molecule kit.

#### RT^2^ Profiler PCR Array

A SYBR green-based quantitative real-time RT^2^ qPCR array was used to profile genes involved in ECM composition as well as adhesion molecule deposition via cell-cell and cell-matrix interactions according to the manufacturer’s instructions (PAMM-013Z; QIAGEN, Hilden, Germany). The 96-well RT^2^ profile plate contained primers for 84 genes of interest, five housekeeping genes, and three negative control wells. RNA extraction was performed as described previously. Briefly, 111 µL cDNA was added to the master mix, and then 25 µL was pipetted into each well. Real-time qPCR was performed on the Bio-Rad CFX 96 (Bio-Rad) using the following cycles: 1 cycle at 95°C for 10 min, 40 cycles at 95°C for 15 s and 60°C for 1 min. A dissociation curve was performed at the end of the program to ensure the amplification of a single product. The results were normalized against the housekeeping gene and analyzed on an EXCEL-based spreadsheet (available on the QIAGEN website: <u>dataanalysis@.qiagen.com</u>). Ct values above 35 were deemed negative and the normalized expression levels of lasered retinas were evaluated relative to non-lasered controls using the 2^−ΔΔCt^ method. A difference > 0.5 or < 0.5 indicated up- or down-regulation, respectively.

#### Statistical analysis

Values were presented as mean ± SD. Statistical analysis was performed using the Student’s t-test for two groups or one-way ANOVA with Bonferroni post hoc test for multiple comparison. Statistically significant differences were denoted by * with a probability value (*P*) of < 0.05, ** with *P* < 0.01, and highly significant differences by *** with *P* < 0.001 and **** with *P* < 0.0001. Graph Pad Prism 9 (GraphPad Software, Boston, MA, USA) was used for all statistical analyses.

## Data Availability Statement

All the data have been included in supplementary.

## Acknowledgments

The authors thanks Anelia Schweri-Olac for outstanding technical assistance. This work was kindly supported by the MIC imaging facility of the Dept. of BioMedical Research, University of Bern, especially by Selina Steiner and Carlos Wotzkow.

## References

[1] Ishikawa, K., Kannan, R. & Hinton, D. R. Molecular mechanisms of subretinal fibrosis in age-related macular degeneration. Exp Eye Res 142, 19–25, doi:10.1016/j.exer.2015.03.009 (2016).

[2] Gabbiani, G. The myofibroblast in wound healing and fibrocontractive diseases. J Pathol 200, 500–503, doi:10.1002/path.1427 (2003).

[3] Van Leeuwen, E. M. et al. A new perspective on lipid research in age-related macular degeneration. Prog Retin Eye Res 67, 56–86, doi:10.1016/j.preteyeres.2018.04.006 (2018).

[4] Litwińska, Z. et al. The Interplay Between Systemic Inflammatory Factors and MicroRNAs in Age-Related Macular Degeneration. Front Aging Neurosci 11, 286, doi:10.3389/fnagi.2019.00286 (2019).

[5] Wooff, Y., Man, S. M., Aggio-Bruce, R., Natoli, R. & Fernando, N. IL-1 Family Members Mediate Cell Death, Inflammation and Angiogenesis in Retinal Degenerative Diseases.

[6] Natoli, R. et al. Microglia-derived IL-1β promotes chemokine expression by Müller cells and RPE in focal retinal degeneration. Mol Neurodegener 12, 31, doi:10.1186/s13024-017-0175-y (2017).

[7] Spindler, J., Zandi, S., Pfister, I. B., Gerhardt, C. & Garweg, J. G. Cytokine profiles in the aqueous humor and serum of patients with dry and treated wet age-related macular degeneration. PLoS One 13, e0203337, doi:10.1371/journal.pone.0203337 (2018).

[8] Miller, J. W. Beyond VEGF-The Weisenfeld Lecture. Invest Ophthalmol Vis Sci 57, 6911–6918, doi:10.1167/iovs.16-21201 (2016).

[9] Viviani, B., Corsini, E., Binaglia, M., Galli, C. L. & Marinovich, M. Reactive oxygen species generated by glia are responsible for neuron death induced by human immunodeficiency virus-glycoprotein 120 in vitro. Neuroscience 107, 51–58, 10.1016/S0306-4522(01)00332-3 (2001).

[10] Viviani, B. et al. Interleukin-1β Enhances NMDA Receptor-Mediated Intracellular Calcium Increase through Activation of the Src Family of Kinases. The Journal of Neuroscience 23, 8692–8700, doi:10.1523/jneurosci.23-25-08692.2003 (2003).

[11] Wang, H., Han, X., Wittchen, E. S. & Hartnett, M. E. TNF-α mediates choroidal neovascularization by upregulating VEGF expression in RPE through ROS-dependent β-catenin activation. Mol Vis 22, 116–128 (2016).

[12] Alshoubaki, Y. K., Nayer, B., Das, S. & Martino, M. M. Modulation of the Activity of Stem and Progenitor Cells by Immune Cells. Stem Cells Transl Med 11, 248–258, doi:10.1093/stcltm/szab022 (2022).

[13] Olivares-González, L., Velasco, S., Campillo, I. & Rodrigo, R. Retinal Inflammation, Cell Death and Inherited Retinal Dystrophies. Int J Mol Sci 22, doi:10.3390/ijms22042096 (2021).

[14] Gao, C. et al. Pirfenidone Alleviates Choroidal Neovascular Fibrosis through TGF-β/Smad Signaling Pathway. Journal of Ophthalmology 2021, 8846708, doi:10.1155/2021/8846708 (2021).

[15] Ma, W. et al. Absence of TGFβ signaling in retinal microglia induces retinal degeneration and exacerbates choroidal neovascularization. eLife 8, e42049, doi:10.7554/eLife.42049 (2019).

[16] Hewitson, T. D. et al. Pirfenidone reduces in vitro rat renal fibroblast activation and mitogenesis. J Nephrol 14, 453–460 (2001).

[17] Ruwanpura, S. M., Thomas, B. J. & Bardin, P. G. Pirfenidone: Molecular Mechanisms and Potential Clinical Applications in Lung Disease. Am J Respir Cell Mol Biol 62, 413–422, doi:10.1165/rcmb.2019-0328TR (2020).

[18] Lopez-de la Mora, D. A., et al. Role and New Insights of Pirfenidone in Fibrotic Diseases. International Journal of Medical Sciences 12, 840–847, doi:10.7150/ijms.11579 (2015).

[19] Meng, X.-m., Nikolic-Paterson, D. J. & Lan, H. Y. TGF-β: the master regulator of fibrosis. Nature Reviews Nephrology 12, 325–338, doi:10.1038/nrneph.2016.48 (2016).

[20] Diaz-Palomera, C. D. et al. Topical Pirfenidone-Loaded Liposomes Ophthalmic Formulation Reduces Haze Development after Corneal Alkali Burn in Mice. Pharmaceutics 14, doi:10.3390/pharmaceutics14020316 (2022).

[21] Khanum, B. N. M. K. et al. Pirfenidone inhibits post-traumatic proliferative vitreoretinopathy. Eye 31, 1317–1328, doi:10.1038/eye.2017.21 (2017).

[22] Talpan, D., Salla, S., Seidelmann, N., Walter, P. & Fuest, M. Antifibrotic Effects of Caffeine, Curcumin and Pirfenidone in Primary Human Keratocytes. International Journal of Molecular Sciences 24, 1461 (2023).

[23] Tracy, L. E., Minasian, R. A. & Caterson, E. J. Extracellular Matrix and Dermal Fibroblast Function in the Healing Wound. Adv Wound Care (New Rochelle) 5, 119–136, doi:10.1089/wound.2014.0561 (2016).

[24] Zandi, S. et al. Animal model of subretinal fibrosis without active choroidal neovascularization. Experimental Eye Research, 109428, 10.1016/j.exer.2023.109428 (2023).

[25] Zhao, Z. et al. TGF-β promotes pericyte-myofibroblast transition in subretinal fibrosis through the Smad2/3 and Akt/mTOR pathways. Experimental & Molecular Medicine 54, 673–684, doi:10.1038/s12276-022-00778-0 (2022).

[26] Shu, D. Y., Butcher, E. & Saint-Geniez, M. EMT and EndMT: Emerging Roles in Age-Related Macular Degeneration. Int J Mol Sci 21, doi:10.3390/ijms21124271 (2020).

[27] Hinz, B. Myofibroblasts. Experimental Eye Research 142, 56–70, 10.1016/j.exer.2015.07.009 (2016).

[28] An, D., Chung-Wah-Cheong, J., Yu, D. Y. & Balaratnasingam, C. Alpha-Smooth Muscle Actin Expression and Parafoveal Blood Flow Pathways Are Altered in Preclinical Diabetic Retinopathy. Invest Ophthalmol Vis Sci 63, 8, doi:10.1167/iovs.63.5.8 (2022).

[29] Eming, S. A., Krieg, T. & Davidson, J. M. Inflammation in Wound Repair: Molecular and Cellular Mechanisms. Journal of Investigative Dermatology 127, 514–525, 10.1038/sj.jid.5700701 (2007).

[30] Damico, F. M., Gasparin, F., Scolari, M. R., Pedral, L. S. & Takahashi, B. S. New approaches and potential treatments for dry age-related macular degeneration. Arq Bras Oftalmol 75, 71–76, doi:10.1590/s0004-27492012000100016 (2012).

[31] Donato, R. et al. S100B’s double life: Intracellular regulator and extracellular signal. Biochimica et Biophysica Acta (BBA) - Molecular Cell Research 1793, 1008–1022, 10.1016/j.bbamcr.2008.11.009 (2009).

[32] Budatha, M., Zhang, J. & Schwartz, M. A. Fibronectin-Mediated Inflammatory Signaling Through Integrin α5 in Vascular Remodeling. J Am Heart Assoc 10, e021160, doi:10.1161/jaha.121.021160 (2021).

[33] Van Der Rest, M. & Garrone, R. Collagen family of proteins. The FASEB Journal 5, 2814–2823, 10.1096/fasebj.5.13.1916105 (1991).

[34] Singh, D., Rai, V. & Agrawal, D. K. Regulation of Collagen I and Collagen III in Tissue Injury and Regeneration. Cardiol Cardiovasc Med 7, 5–16, doi:10.26502/fccm.92920302 (2023).

[35] Halfter, W., Willem, M. & Mayer, U. Basement membrane-dependent survival of retinal ganglion cells. Invest Ophthalmol Vis Sci 46, 1000–1009, doi:10.1167/iovs.04-1185 (2005).

[36] Tomino, Y. et al. Asian multicenter trials on urinary type IV collagen in patients with diabetic nephropathy. J Clin Lab Anal 15, 188–192, doi:10.1002/jcla.1026 (2001).

[37] Urushiyama, H. et al. Role of α1 and α2 chains of type IV collagen in early fibrotic lesions of idiopathic interstitial pneumonias and migration of lung fibroblasts. Laboratory Investigation 95, 872–885, doi:10.1038/labinvest.2015.66 (2015).

[38] Ge, Q. et al. Fibulin1C peptide induces cell attachment and extracellular matrix deposition in lung fibroblasts. Scientific Reports 5, 9496, doi:10.1038/srep09496 (2015).

[39] Wight, T. N. et al. Versican-A Critical Extracellular Matrix Regulator of Immunity and Inflammation. Front Immunol 11, 512, doi:10.3389/fimmu.2020.00512 (2020).

[40] Song, I. & Dityatev, A. Crosstalk between glia, extracellular matrix and neurons. Brain Research Bulletin 136, 101-108, 10.1016/j.brainresbull.2017.03.003 (2018).

[41] Kawashima, H. et al. Binding of a large chondroitin sulfate/dermatan sulfate proteoglycan, versican, to L-selectin, P-selectin, and CD44. *J Biol Chem* **275**, 35448-35456, doi:10.1074/jbc.M003387200 (2000).

[42] Zheng, P.-S. et al. PG-M/versican binds to P-selectin glycoprotein ligand-1 and mediates leukocyte aggregation. Journal of Cell Science 117, 5887–5895, doi:10.1242/jcs.01516 (2004).

[43] Wu, Y. J., Pierre, D. P. L. A., Wu, J., Yee, A. J. & Yang, B. B. The interaction of versican with its binding partners. Cell Research 15, 483–494, doi:10.1038/sj.cr.7290318 (2005).

[44] Hirano, Y., Sakurai, E., Matsubara, A. & Ogura, Y. Suppression of ICAM-1 in Retinal and Choroidal Endothelial Cells by Plasmid Small-Interfering RNAs In Vivo. Investigative Ophthalmology & Visual Science 51, 508–515, doi:10.1167/iovs.09-3457 (2010).

[45] Granot, E., Shouval, D. & Ashur, Y. Cell adhesion molecules and hyaluronic acid as markers of inflammation, fibrosis and response to antiviral therapy in chronic hepatitis C patients. Mediators of Inflammation 10, 274192, doi:10.1080/09629350120093722 (2001).

[46] Yang, X., Chung, J.-Y., Rai, U. & Esumi, N. Cadherins in the retinal pigment epithelium (RPE) revisited: P-cadherin is the highly dominant cadherin expressed in human and mouse RPE in vivo. PLOS ONE 13, e0191279, doi:10.1371/journal.pone.0191279 (2018).

[47] Polisetti, N., Sharaf, L., Martin, G., Schlunck, G. & Reinhard, T. P-Cadherin Is Expressed by Epithelial Progenitor Cells and Melanocytes in the Human Corneal Limbus. Cells 11, 1975 (2022).

[48] Aimo, A. et al. Pirfenidone for Idiopathic Pulmonary Fibrosis and Beyond. Card Fail Rev 8, e12, doi:10.15420/cfr.2021.30 (2022).

[49] Adler, M. et al. Principles of Cell Circuits for Tissue Repair and Fibrosis. iScience 23, 100841, doi:10.1016/j.isci.2020.100841 (2020).

[50] Jun, J. I. & Lau, L. F. Resolution of organ fibrosis. J Clin Invest 128, 97–107, doi:10.1172/jci93563 (2018).

[51] Rodrigues, R. M. et al. E-Selectin-Dependent Inflammation and Lipolysis in Adipose Tissue Exacerbate Steatosis-to-NASH Progression via S100A8/9. Cellular and Molecular Gastroenterology and Hepatology 13, 151–171, 10.1016/j.jcmgh.2021.08.002 (2022).

[52] Zhang, N., Liu, Z., Yao, L., Mehta-D’souza, P. & McEver, R. P. P-Selectin Expressed by a Human SELP Transgene Is Atherogenic in Apolipoprotein E-Deficient Mice. Arterioscler Thromb Vasc Biol 36, 1114–1121, doi:10.1161/atvbaha.116.307437 (2016).

[53] Corona, A. & Blobe, G. C. The role of the extracellular matrix protein TGFBI in cancer. Cellular Signalling 84, 110028, 10.1016/j.cellsig.2021.110028 (2021).

