## Supplemental figures for "Modulation of Extracellular Matrix Composition and Chronic Inflammation by Pirfenidone Promotes Scar Reduction in Retinal Wound Repair"

**Supplementary materials**

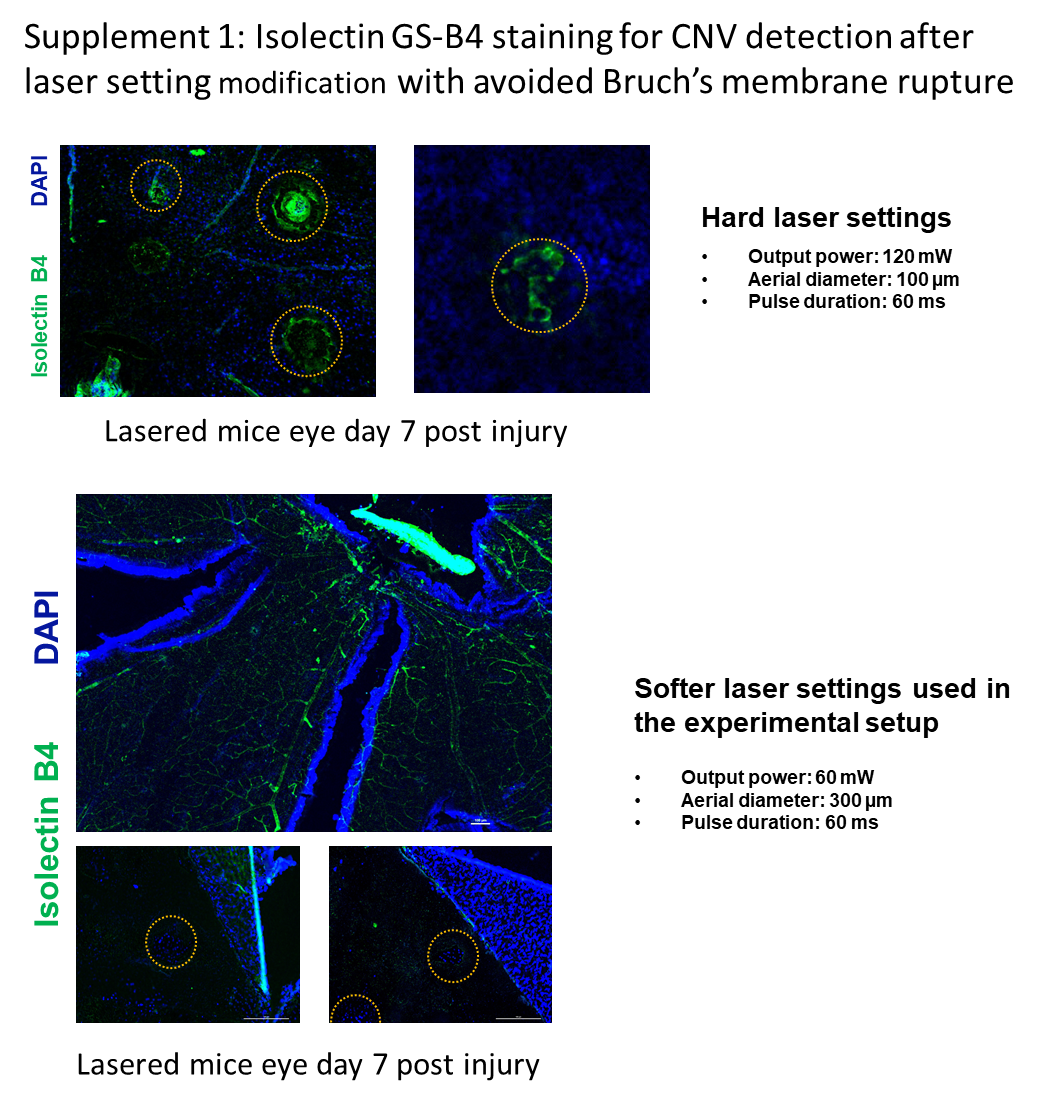

Supplementary Figure 1: Isolectin GS-B4 staining for CNV detection without Bruch's membrane rupture after modification of the laser setting.

Supplementary Table 1: Fold change of ECM component proteins post lasered samples (significant changes in bold)

|  | **Day 7** | | **D21** | | **D35** | | **D49** | |
| --- | --- | --- | --- | --- | --- | --- | --- | --- |
| **Gene Symbol** | **Fold Change** | **P-Value** | **Fold Change** | **P-Value** | **Fold Change** | **P-Value** | **Fold Change** | **P-Value** |
| *Col1a1* | 2.93 | 0.26 | 0.41 | 0.40 | 1.02 | 0.55 | 1.21 | 0.73 |
| *Col2a1* | 1.68 | 0.52 | 0.91 | 0.90 | 1.96 | 0.31 | 2.31 | 0.18 |
| *Col3a1* | 1.91 | 0.28 | 0.4 | 0.13 | 0.32 | 0.07 | 0.35 | 0.07 |
| *Col4a1* | 0.58 | 0.12 | 1.11 | 0.62 | 0.72 | 0.17 | 0.74 | 0.21 |
| *Col4a2* | 0.99 | 0.80 | 0.77 | 0.88 | 1.04 | 0.81 | 1.28 | 0.81 |
| *Col4a3* | 2.54 | 0.08 | 1.5 | 0.34 | **3.15** | 0.00 | **4.32** | 0.01 |
| *Col5a1* | 2.26 | 0.41 | 0.87 | 0.93 | 2.16 | 0.44 | 2.85 | 0.17 |
| *Col6a1* | 1.46 | 0.92 | 0.56 | 0.51 | 1.33 | 0.86 | 1.69 | 0.77 |
| *Ecm1* | 1.84 | 0.49 | 0.94 | 0.87 | 1.81 | 0.49 | 2.42 | 0.18 |
| *Emilin1* | 2.02 | 0.55 | 0.89 | 0.90 | 1.72 | 0.82 | 2.23 | 0.41 |
| *Fbln1* | 4.89 | 0.22 | 5.45 | 0.13 | **12.38** | 0.00 | **4.35** | 0.00 |
| *Fn1* | 1.53 | 0.81 | 0.6 | 0.55 | 1.36 | 0.94 | 1.82 | 0.58 |
| *Hapln1* | 1.29 | 0.66 | 0.7 | 0.41 | 1.12 | 0.96 | 1.31 | 0.65 |
| *Lama1* | 1.72 | 0.67 | 0.8 | 0.65 | 1.56 | 0.85 | 2.15 | 0.37 |
| *Lama2* | 1.27 | 0.51 | 0.98 | 0.92 | 0.91 | 0.67 | 1.06 | 0.95 |
| *Lama3* | 1.42 | 0.46 | 1.03 | 0.78 | 1.27 | 0.64 | 1.74 | 0.18 |
| *Lamb2* | 1.71 | 0.39 | 1.08 | 0.98 | 1.36 | 0.74 | 1.59 | 0.47 |
| *Lamb3* | 1.26 | 0.89 | 0.7 | 0.44 | 1.11 | 0.84 | 1.67 | 0.49 |
| *Lamc1* | 2.34 | 0.66 | 0.62 | 0.67 | 2.44 | 0.62 | 3.47 | 0.24 |
| *Sparc* | 1.46 | 0.82 | 0.88 | 0.77 | 1.11 | 0.72 | 1.27 | 0.92 |
| *Spock1* | 1.38 | 0.91 | 0.73 | 0.64 | 1.54 | 0.78 | 1.87 | 0.49 |
| *Spp1* | 1.08 | 0.81 | 0.85 | 0.56 | 1.11 | 0.78 | 1.16 | 0.65 |
| *Syt1* | 1.52 | 0.63 | 0.6 | 0.54 | 1.5 | 0.67 | 1.95 | 0.29 |
| *Tnc* | 1.98 | 0.72 | 0.67 | 0.66 | 1.64 | 1.00 | 2.56 | 0.38 |
| *Vcan* | 2.09 | 0.26 | 2.03 | 0.22 | **3.76** | 0.00 | **2.34** | 0.03 |
| *Vtn* | 1.56 | 0.51 | 0.78 | 0.72 | 1.52 | 0.54 | 2.03 | 0.17 |

Supplementary Table 2: Transmembrane and adhesion molecules fold change laser treated samples (significant changes in bold)

|  | **Day 7** | | **D21** | | **D35** | | **D49** | |
| --- | --- | --- | --- | --- | --- | --- | --- | --- |
| **Gene Symbol** | **Fold Change** | **P-Value** | **Fold Change** | **P-Value** | **Fold Change** | **P-Value** | **Fold Change** | **P-Value** |
| *Cd44* | 1.75 | 0.30 | 1.18 | 0.63 | 1.79 | 0.15 | 1.98 | 0.11 |
| *Cdh1* | 1.51 | 0.35 | 2.33 | 0.36 | 1.61 | 0.23 | 1.84 | 0.15 |
| *Cdh2* | 2.96 | 0.65 | 0.66 | 0.63 | 2.99 | 0.68 | 3.84 | 0.35 |
| *Cdh3* | **0.68** | 0.03 | 0.81 | 0.23 | **0.65** | 0.01 | 1.03 | 0.77 |
| *Cdh4* | 1.82 | 0.70 | 0.6 | 0.66 | 1.91 | 0.65 | 2.59 | 0.25 |
| *Cntn1* | 1.29 | 0.61 | 0.63 | 0.48 | 1.29 | 0.59 | 1.5 | 0.29 |
| *Ctnna1* | 1.66 | 0.54 | 0.75 | 0.76 | 1.59 | 0.61 | 1.83 | 0.39 |
| *Ctnna2* | 1.34 | 0.76 | 0.65 | 0.59 | 1.27 | 0.89 | 1.6 | 0.47 |
| *Ctnnb1* | 1.52 | 0.55 | 0.72 | 0.60 | 1.36 | 0.75 | 1.73 | 0.35 |
| *Icam1* | **3.01** | 0.05 | 1.35 | 0.56 | 2.53 | 0.09 | 3.59 | 0.06 |
| *Itga2* | 1.34 | 0.47 | 0.62 | 0.54 | 1.06 | 0.99 | 1.64 | 0.12 |
| *Itga3* | 1.13 | 0.81 | 0.74 | 0.60 | 0.95 | 0.89 | 1.44 | 0.36 |
| *Itga4* | 1.29 | 0.51 | 0.71 | 0.40 | 1.06 | 0.93 | 1.56 | 0.12 |
| *Itga5* | 2.04 | 0.66 | 0.7 | 0.70 | 2.04 | 0.68 | 3.06 | 0.24 |
| *Itgae* | 1.54 | 0.53 | 1.47 | 0.60 | 1.58 | 0.48 | 2.36 | 0.07 |
| *Itgal* | 2.08 | 0.11 | 1.2 | 0.69 | 1.89 | 0.08 | **3.69** | 0.03 |
| *Itgam* | 2.01 | 0.24 | 0.82 | 0.86 | 1.35 | 0.59 | 1.71 | 0.26 |
| *Itgav* | 1.23 | 0.60 | 0.67 | 0.50 | 1.27 | 0.47 | 1.59 | 0.15 |
| *Itgax* | 1.27 | 0.46 | 1.57 | 0.23 | **0.65** | 0.01 | 1.28 | 0.27 |
| *Itgb1* | 1.36 | 0.63 | 0.69 | 0.65 | 1.3 | 0.74 | 1.52 | 0.45 |
| *Itgb2* | 2.34 | 0.16 | 0.84 | 0.95 | 1.84 | 0.15 | 2.11 | 0.11 |
| *Itgb3* | 1.43 | 0.76 | 0.65 | 0.65 | 0.93 | 0.56 | 1.65 | 0.52 |
| *Itgb4* | 2.07 | 0.68 | 0.73 | 0.65 | 2.2 | 0.59 | 2.83 | 0.28 |
| *Ncam1* | 1.91 | 0.50 | 0.73 | 0.71 | 1.94 | 0.47 | 2.53 | 0.19 |
| *Ncam2* | 1.08 | 0.87 | 0.57 | 0.36 | 1.07 | 1.00 | 1.28 | 0.54 |
| *Pecam1* | 1.63 | 0.56 | 0.54 | 0.41 | 1.38 | 0.86 | 2 | 0.29 |
| *Postn* | 1.48 | 0.47 | 0.54 | 0.29 | 0.8 | 0.45 | 1.01 | 0.82 |
| *Sele* | 0.95 | 0.98 | 1.15 | 0.39 | **0.59** | 0.00 | 1.06 | 0.48 |
| *Sell* | **0.7** | 0.01 | 0.88 | 0.68 | **0.62** | 0.00 | 1.15 | 0.52 |
| *Selp* | 1.5 | 0.29 | 1.14 | 0.30 | 1.13 | 0.34 | **2.23** | 0.03 |
| *Sgce* | 1.27 | 0.71 | 0.8 | 0.69 | 1.35 | 0.64 | 1.43 | 0.53 |
| *Thbs1* | 1.29 | 0.92 | 0.62 | 0.57 | 1.48 | 0.72 | 1.22 | 0.92 |
| *Thbs2* | 2.18 | 0.17 | 1.02 | 0.75 | 1.71 | 0.08 | 1.67 | 0.16 |
| *Thbs3* | 2.13 | 0.56 | 0.8 | 0.79 | 1.86 | 0.76 | 2.57 | 0.33 |
| *Vcam1* | 1.52 | 0.50 | 0.71 | 0.57 | 1.35 | 0.71 | 1.61 | 0.39 |

Supplementary Table 3: Fold change data of ECM protease and inhibitors post laser (significant changes in bold)

|  | **Day 7** | | **D21** | | **D35** | | **D49** | |
| --- | --- | --- | --- | --- | --- | --- | --- | --- |
| **Gene Symbol** | **Fold Change** | **P-Value** | **Fold Change** | **P-Value** | **Fold Change** | **P-Value** | **Fold Change** | **P-Value** |
| *Adamts1* | 2 | 0.23 | 0.99 | 0.95 | 1.6 | 0.41 | 1.39 | 0.66 |
| *Adamts2* | 1.44 | 0.40 | 0.73 | 0.43 | 0.97 | 0.85 | 1 | 0.88 |
| *Adamts5* | 1.61 | 0.31 | 0.93 | 0.93 | 1.27 | 0.52 | 1.68 | 0.11 |
| *Adamts8* | 1.5 | 0.90 | 0.73 | 0.56 | 1.49 | 0.95 | 2.17 | 0.38 |
| *Mmp10* | 0.77 | 0.08 | 0.9 | 0.74 | **0.65** | 0.01 | 1.34 | 0.30 |
| *Mmp11* | 1.67 | 0.51 | 0.69 | 0.90 | 1.77 | 0.40 | 2.34 | 0.16 |
| *Mmp12* | 0.97 | 0.85 | 0.94 | 0.89 | 0.88 | 0.54 | 1.44 | 0.36 |
| *Mmp13* | 0.97 | 0.94 | 1.22 | 0.47 | 0.96 | 0.99 | 1.62 | 0.12 |
| *Mmp14* | 2.05 | 0.74 | 0.71 | 0.66 | 2.25 | 0.62 | 2.65 | 0.40 |
| *Mmp15* | 2.17 | 0.74 | 0.62 | 0.65 | 2.03 | 0.87 | 2.75 | 0.45 |
| *Mmp1a* | 1.31 | 0.79 | 0.56 | 0.48 | 1.23 | 0.98 | 1.55 | 0.54 |
| *Mmp2* | 1.59 | 0.63 | 0.64 | 0.58 | 1.51 | 0.73 | 1.89 | 0.38 |
| *Mmp3* | 1.72 | 0.22 | 0.87 | 0.20 | **0.79** | 0.01 | 1.46 | 0.09 |
| *Mmp7* | 0.82 | 0.29 | 0.81 | 0.23 | **0.65** | 0.01 | 1.07 | 0.60 |
| *Mmp8* | **0.75** | 0.05 | 0.9 | 0.68 | **0.65** | 0.01 | 1.32 | 0.37 |
| *Mmp9* | 1.51 | 0.87 | 0.69 | 0.57 | 1.45 | 1.00 | 1.85 | 0.61 |
| *Timp1* | 2.06 | 0.35 | 0.8 | 0.73 | 1.51 | 0.79 | 1.91 | 0.41 |
| *Timp2* | 1.86 | 0.51 | 0.87 | 0.82 | 1.84 | 0.52 | 2.3 | 0.25 |
| *Timp3* | 1.34 | 0.68 | 0.8 | 0.82 | 1.55 | 0.45 | 1.54 | 0.46 |

Supplementary Table 4: Fold change of other ECM proteins post laser (significant changes in bold)

|  | **Day 7** | | **D21** | | **D35** | | **D49** | |
| --- | --- | --- | --- | --- | --- | --- | --- | --- |
| **Gene Symbol** | **Fold Change** | **P-Value** | **Fold Change** | **P-Value** | **Fold Change** | **P-Value** | **Fold Change** | **P-Value** |
| *Entpd1* | 1.77 | 0.55 | 0.78 | 0.78 | 1.26 | 0.97 | 2.02 | 0.36 |
| *Hc* | 0.87 | 0.67 | 0.81 | 0.23 | **0.65** | 0.01 | 1.14 | 0.11 |
| *Tgfbi* | 3.46 | 0.06 | 1.7 | 0.37 | 2.27 | 0.11 | 1.97 | 0.20 |

**Supplementary Table 5: Fold change of ECM components PFD samples** (significant changes in bold)

|  | **Day 21 PFD** | | **D35 PFD** | | **D42 PFD** | | | | **D42 PFD D35** | |
| --- | --- | --- | --- | --- | --- | --- | --- | --- | --- | --- |
| **Gene Symbol** | **Fold Change** | **P-Value** | **Fold Change** | **P-Value** | | **Fold Change** | **P-Value** | **Fold Change** | | **P-Value** |
| *Cd44* | 1.17 | 0.98 | 1.13 | 0.90 | | 1.27 | 0.83 | 0.98 | | 0.66 |
| *Cdh1* | 0.81 | 0.70 | 0.81 | 0.63 | | 0.98 | 0.83 | 0.67 | | 0.17 |
| *Cdh2* | 1.68 | 0.54 | 1.81 | 0.62 | | 1.36 | 0.40 | 1.31 | | 0.38 |
| *Cdh3* | 0.67 | 0.28 | 0.66 | 0.26 | | 0.5 | 0.13 | 0.45 | | 0.10 |
| *Cdh4* | 1.1 | 0.50 | 1.28 | 0.63 | | 1.13 | 0.52 | 1.09 | | 0.49 |
| *Cntn1* | 0.73 | 0.30 | 0.83 | 0.46 | | 0.79 | 0.37 | 0.66 | | 0.25 |
| *Ctnna1* | 0.94 | 0.49 | 1.03 | 0.63 | | 0.98 | 0.54 | 0.8 | | 0.37 |
| *Ctnna2* | 0.79 | 0.37 | **0.89** | 0.50 | | 0.81 | 0.39 | 0.74 | | 0.33 |
| *Ctnnb1* | 0.89 | 0.48 | 1.03 | 0.70 | | 0.81 | 0.40 | 0.75 | | 0.34 |
| *Icam1* | 2.64 | 0.09 | **2.8** | 0.05 | | 1.96 | 0.29 | 2.3 | | 0.14 |
| *Itga2* | 1.13 | 0.86 | 1.07 | 0.98 | | 1.12 | 0.91 | 1.13 | | 0.87 |
| *Itga3* | 0.7 | 0.27 | 0.79 | 0.36 | | 0.74 | 0.30 | 0.69 | | 0.24 |
| *Itga4* | 0.64 | 0.15 | 0.78 | 0.30 | | 0.8 | 0.35 | 0.82 | | 0.37 |
| *Itga5* | 1.19 | 0.52 | 1.48 | 0.76 | | 1.37 | 0.64 | 1 | | 0.39 |
| *Itgae* | 2 | 0.20 | 27.33 | 0.37 | | 1.84 | 0.26 | 1.9 | | 0.21 |
| *Itgal* | 1.23 | 0.93 | 1.43 | 0.81 | | 1.19 | 0.90 | 1.04 | | 0.73 |
| *Itgam* | 1.09 | 0.90 | 1.04 | 0.75 | | 0.97 | 0.61 | 0.9 | | 0.51 |
| *Itgav* | 0.77 | 0.32 | 0.82 | 0.42 | | 0.82 | 0.39 | 0.72 | | 0.26 |
| *Itgax* | 0.87 | 0.57 | 0.72 | 0.38 | | 0.59 | 0.18 | 0.66 | | 0.27 |
| *Itgb1* | 0.82 | 0.42 | 0.91 | 0.55 | | 0.84 | 0.43 | 0.82 | | 0.41 |
| *Itgb2* | **2.24** | 0.05 | 2.34 | 0.06 | | **2.38** | 0.02 | **2.27** | | 0.02 |
| *Itgb3* | **4.5** | 0.03 | 3.98 | 0.06 | | **4.02** | 0.02 | **3.77** | | 0.03 |
| *Itgb4* | 1.13 | 0.50 | 1.27 | 0.54 | | 1.35 | 0.61 | 1.13 | | 0.45 |
| *Ncam1* | 1.13 | 0.62 | 1.32 | 0.83 | | 1.19 | 0.67 | 1.11 | | 0.58 |
| *Ncam2* | 0.59 | 0.16 | 0.77 | 0.34 | | 0.7 | 0.24 | 0.67 | | 0.21 |
| *Pecam1* | 0.82 | 0.39 | 0.88 | 0.43 | | 0.76 | 0.34 | 0.84 | | 0.40 |
| *Postn* | 0.44 | 0.13 | 0.54 | 0.17 | | 0.45 | 0.13 | 0.43 | | 0.12 |
| *Sele* | **3.66** | 0.02 | **3.71** | 0.05 | | **3.49** | 0.01 | **3.23** | | 0.00 |
| *Sell* | 2.2 | 0.07 | 2.46 | 0.09 | | **2.29** | 0.01 | **2.38** | | 0.04 |
| *Selp* | 3.93 | 0.01 | **3.27** | 0.01 | | **3.35** | 0.02 | **2.76** | | 0.01 |
| *Sgce* | **0.62** | 0.22 | 0.73 | 0.35 | | 0.61 | 0.21 | 0.56 | | 0.18 |
| *Thbs1* | 1.04 | 0.61 | 0.83 | 0.40 | | 0.83 | 0.39 | 0.66 | | 0.27 |
| *Thbs2* | 1 | 0.86 | 0.94 | 0.65 | | 0.74 | 0.29 | 0.94 | | 0.64 |
| *Thbs3* | 1.56 | 0.91 | 1.7 | 0.96 | | 1.51 | 0.85 | 1.51 | | 0.85 |

Supplementary Table 6: Fold change of Transmembrane and adhesion molecules PFD samples (significant changes in bold)

|  | **Day 21 PFD** | | **D35 PFD** | | **D42 PFD** | | **D42 PFD D35** | |
| --- | --- | --- | --- | --- | --- | --- | --- | --- |
| **Gene Symbol** | **Fold Change** | **P-Value** | **Fold Change** | **P-Value** | **Fold Change** | **P-Value** | **Fold Change** | **P-Value** |
| *Col1a1* | 1.88 | 0.66 | 1.78 | 0.75 | 1.81 | 0.74 | 1.75 | 0.81 |
| *Col2a1* | 1.35 | 0.87 | 1.52 | 0.61 | 1.21 | 0.91 | 1.02 | 0.64 |
| *Col3a1* | 0.39 | 0.04 | **0.29** | 0.02 | **0.32** | 0.02 | **0.28** | 0.02 |
| *Col4a1* | **0.4** | 0.0001 | **0.42** | 0.0010 | **0.39** | 0.0001 | **0.35** | 0.0001 |
| *Col4a2* | 0.48 | 0.17 | 0.66 | 0.26 | 0.57 | 0.21 | 0.54 | 0.19 |
| *Col4a3* | 1.92 | 0.12 | 2.03 | 0.10 | **2.47** | 0.05 | **2.74** | 0.01 |
| *Col5a1* | 1.11 | 0.55 | 1.34 | 0.78 | 1.18 | 0.61 | 1.08 | 0.52 |
| *Col6a1* | 1.15 | 0.59 | 1.28 | 0.71 | 1.06 | 0.51 | 1.05 | 0.50 |
| *Ecm1* | 1.15 | 0.71 | 1.17 | 0.75 | 1.02 | 0.56 | 0.94 | 0.48 |
| *Emilin1* | 1.5 | 0.86 | 1.46 | 0.86 | 1.46 | 0.82 | 1.51 | 0.87 |
| *Fn1* | 0.94 | 0.44 | 1.04 | 0.55 | 0.87 | 0.38 | 0.77 | 0.32 |
| *Hapln1* | 0.61 | 0.28 | 0.84 | 0.48 | 0.7 | 0.32 | 0.67 | 0.29 |
| *Lama1* | 1.01 | 0.48 | 1.13 | 0.60 | 0.84 | 0.37 | 0.89 | 0.39 |
| *Lama2* | 0.5 | 0.09 | 0.52 | 0.07 | **0.43** | 0.05 | 0.52 | 0.06 |
| *Lama3* | 0.87 | 0.50 | 1.03 | 0.84 | 0.8 | 0.38 | 0.87 | 0.50 |
| *Lamb2* | 1.35 | 0.79 | 1.4 | 0.72 | 1.14 | 0.86 | 1.07 | 0.73 |
| *Lamb3* | 2.81 | 0.10 | 2.46 | 0.16 | 2.11 | 0.27 | 2.1 | 0.26 |
| *Lamc1* | 1.47 | 0.66 | 1.65 | 0.78 | 1.42 | 0.61 | 1.33 | 0.55 |
| *Sparc* | 0.74 | 0.32 | 0.73 | 0.35 | 0.57 | 0.23 | 0.54 | 0.21 |
| *Spock1* | 0.68 | 0.28 | 0.85 | 0.44 | 0.78 | 0.35 | 0.69 | 0.29 |
| *Syt1* | 0.87 | 0.46 | 1.09 | 0.73 | 1.04 | 0.66 | 1.02 | 0.62 |
| *Tnc* | 1.03 | 0.42 | 1.38 | 0.69 | 1.14 | 0.47 | 0.89 | 0.33 |
| *Vcan* | 0.88 | 0.98 | 0.65 | 0.06 | 0.75 | 0.88 | 0.87 | 0.99 |
| *Vtn* | 1.07 | 0.73 | 1.17 | 0.97 | 0.9 | 0.51 | 1.62 | 0.47 |

Supplementary Table 7: Fold change of ECM protease and inhibitors PFD samples (significant changes in bold)

|  | **Day 21 PFD** | | **D35 PFD** | | **D42 PFD** | | **D42 PFD D35** | |
| --- | --- | --- | --- | --- | --- | --- | --- | --- |
| **Gene Symbol** | **Fold Change** | **P-Value** | **Fold Change** | **P-Value** | **Fold Change** | **P-Value** | **Fold Change** | **P-Value** |
| *Adamts1* | 0.87 | 0.46 | 1.11 | 0.82 | 0.81 | 0.41 | 0.67 | 0.27 |
| *Adamts2* | 1.36 | 0.51 | 1.17 | 0.84 | 1.34 | 0.50 | 0.96 | 0.68 |
| *Adamts5* | 0.88 | 0.51 | 0.92 | 0.61 | 0.8 | 0.36 | 0.75 | 0.29 |
| *Adamts8* | 1.95 | 0.55 | 2.28 | 0.31 | 2.07 | 0.46 | 1.77 | 0.72 |
| *Mmp10* | 1.51 | 0.30 | **2.25** | 0.04 | 1.52 | 0.22 | 1.51 | 0.24 |
| *Mmp11* | 1.18 | 0.83 | 1.11 | 0.70 | 1.18 | 0.81 | 1.01 | 0.57 |
| *Mmp12* | 2.07 | 0.10 | **2.16** | 0.05 | 2.18 | 0.06 | **2.09** | 0.04 |
| *Mmp13* | 1.6 | 0.11 | 1.83 | 0.17 | 1.28 | 0.33 | 1.32 | 0.26 |
| *Mmp14* | 0.78 | 0.28 | 0.83 | 0.31 | 0.93 | 0.34 | 0.64 | 0.23 |
| *Mmp15* | 1.21 | 0.48 | 1.37 | 0.56 | 1.32 | 0.53 | 1.07 | 0.39 |
| *Mmp1a* | 0.71 | 0.31 | 0.92 | 0.51 | 0.87 | 0.46 | 0.82 | 0.40 |
| *Mmp2* | 1.09 | 0.65 | 1.15 | 0.74 | 1.12 | 0.67 | 0.88 | 0.43 |
| *Mmp3* | 1.1 | 0.84 | 0.88 | 0.77 | 0.63 | 0.25 | 0.6 | 0.24 |
| *Mmp7* | 1.98 | 0.11 | **2.43** | 0.03 | 1.67 | 0.15 | **1.92** | 0.05 |
| *Mmp8* | **3.17** | 0.04 | **3.56** | 0.03 | **3.26** | 0.00 | **3.03** | 0.01 |
| *Mmp9* | 0.82 | 0.36 | 1.12 | 0.59 | 0.94 | 0.43 | 0.87 | 0.38 |
| *Timp1* | 2.69 | 0.20 | 3.16 | 0.10 | 3.07 | 0.10 | 2.4 | 0.27 |
| *Timp2* | 0.97 | 0.50 | 1.12 | 0.65 | 0.95 | 0.47 | 0.93 | 0.46 |
| *Timp3* | 0.81 | 0.39 | 0.76 | 0.38 | 0.76 | 0.34 | 0.67 | 0.27 |

Supplementary Table 8: Fold change of other ECM proteins PFD samples (significant changes in bold)

|  | **Day 21 PFD** | | **D35 PFD** | | **D42 PFD** | | **D42 PFD D35** | |
| --- | --- | --- | --- | --- | --- | --- | --- | --- |
| **Gene Symbol** | **Fold Change** | **P-Value** | **Fold Change** | **P-Value** | **Fold Change** | **P-Value** | **Fold Change** | **P-Value** |
| *Ctgf* | 0.55 | 0.14 | 0.86 | 0.49 | 0.6 | 0.15 | 0.64 | 0.18 |
| *Entpd1* | 1.69 | 0.66 | 1.67 | 0.69 | 1.69 | 0.67 | 1.53 | 0.86 |
| *Hc* | 1.02 | 0.96 | 1.26 | 0.57 | 1.04 | 0.98 | 1.01 | 0.86 |
| *Tgfbi* | **2.9** | 0.04 | **3.16** | 0.04 | **3.01** | 0.02 | **2.7** | 0.03 |
